# Mitonuclear interactions shape male cuticular hydrocarbon profiles with consequences on mating success

**DOI:** 10.64898/2026.03.31.715324

**Authors:** Tom M. Allison, Stuart Harrison, Nick Lane, M. Florencia Camus

## Abstract

Sexual signals are thought to reflect metabolic capacity, allowing females to assess male genetic quality. In insects, cuticular hydrocarbons (CHCs) are central to mate recognition and sexual signalling, and their biosynthesis is directly tied to mitochondrial metabolism. Because mitochondrial performance requires coordination between the mitochondrial and nuclear genomes, non-compatible genomes may disrupt CHC production and reduce male attractiveness. We tested this prediction using a global *Drosophila melanogaster* mitonuclear panel comprising 80 cybrid genotypes. Multivariate analyses of male CHC profiles revealed strong nuclear effects, smaller but significant mitochondrial effects, and substantial non-additive mitonuclear interactions that accounted for ~10% of the variance after controlling for body mass. These interactions reorganised CHC blends in genotype-specific ways, with certain hydrocarbons contributing disproportionately to differentiation. In behavioural assays, females preferentially mated with males whose mitonuclear genomes were coadapted. Conversely, coadapted males had higher copulation success than males presenting disrupted combinations to the female. Our results demonstrate that mitonuclear compatibility influences the production of sexual signals and shapes reproductive outcomes, linking genomic interactions to mate choice.

## 1. Introduction

Sexual signals are some of the most elaborate and costly traits in nature, ranging from bird songs and courtship dances to the colour patterns and chemical displays of insects. A central question in evolutionary biology is why such traits remain honest indicators of quality in the face of strong sexual selection. Theory predicts that sexual signals can be maintained if their expression is condition-dependent and therefore costly to produce [1–4]. Under this framework, only individuals in good physiological condition can sustain the energetic costs of developing or displaying an attractive signal, while low-quality individuals are constrained to produce less elaborate displays. Much empirical support for condition-dependent signalling has come from systems where resource acquisition is variable [5], or where signals are tied to immune function [6, 7], or oxidative stress [8, 9]. However, an underexplored link is the role of energy metabolism itself. Metabolic efficiency is a fundamental determinant of condition because it underpins growth, survival, and reproductive effort. Individuals with a greater capacity to generate usable energy should be better able to allocate resources to energetically demanding traits such as fighting ability, ornament maintenance, or sustained courtship performance. This leads to the prediction that sexual signals could be readouts of metabolic capacity, providing females with reliable information about a male’s physiological quality.

In insects, cuticular hydrocarbons (CHCs) provide a powerful model to test this idea. CHCs are long-chain hydrocarbons that coat the cuticle, serving both to prevent desiccation and as chemicals mediating mate recognition, sexual signalling [10–13], and sometimes species isolation [10, 14]. CHC blends are highly diverse, often species-specific, and their quantitative composition can influence female mate choice [15]. Importantly, CHC biosynthesis depends on fatty-acid precursors derived from acetyl-CoA and the Krebs cycle [16]. This direct metabolic link suggests that variation in mitochondrial function could translate into differences in CHC production, making them ideal candidate traits for testing the hypothesis that sexual signals honestly reflect metabolic efficiency.

Mitochondria are central to energy metabolism, producing ATP and biosynthetic intermediates via oxidative phosphorylation [17]. They are unusual among cellular components in retaining their own genome (mtDNA), which encodes essential subunits of the OXPHOS complexes [18]. Most mitochondrial proteins are however nuclear encoded. As a result, efficient mitochondrial function depends on the tight coordination of gene products from both genomes [19]. Disruption of this coordination (through hybridisation, introgression, or laboratory construction of cytoplasmic hybrids) can lead to mitonuclear incompatibilities that reduce energy efficiency. Such incompatibilities have been linked to reduced fertility, shortened lifespan, slower development, and impaired stress tolerance in diverse taxa [19–21].

Despite growing recognition of their role in organismal fitness, the potential influence of mitonuclear interactions on sexually selected traits has received little attention [22]. This gap is surprising given that sexual selection could act as a powerful force reinforcing mitonuclear coadaptation. If females preferentially mate with males who signal superior metabolic efficiency, and if such efficiency depends on coadapted mitonuclear combinations within the male, then mate choice may bias reproduction toward genomically compatible pairings. Conversely, males with disrupted genomes and impaired mitochondrial function may produce weaker or altered sexual signals, resulting in reduced mating success and direct sexual selection against mitonuclear incompatibility. *Drosophila* cuticular hydrocarbons offer an ideal system in which to explore these ideas. CHCs are well-characterised sexual signals known to influence mate choice, and their production is metabolically costly [23]. Moreover, *Drosophila* are amenable to experimental manipulation of mitonuclear genotypes via the construction of cytoplasmic hybrid (cybrid) panels. These panels permit independent assortment of mitochondrial haplotypes and nuclear backgrounds, allowing the relative contributions of each genome, and their interaction, to be directly quantified.

Here we use a global *Drosophila melanogaster* mitonuclear panel comprising 81 genotypes to test whether mitonuclear interactions shape the production of male CHCs and whether females discriminate among males on this basis. Specifically, we ask three questions. First, do mitonuclear interactions structure CHC profiles beyond the additive effects of the nuclear and mitochondrial genomes alone? Second, which individual hydrocarbons contribute most strongly to differentiation among mitonuclear genotypes, and do they align with axes of mito– nuclear divergence? Third, do these differences in CHC composition have behavioural consequences, such that females preferentially mate with males whose mitonuclear genome is coadapted, or whose mitonuclear background is compatible with their own? By integrating multivariate analyses of pheromone architecture with behavioural assays of mate choice, our study provides experimental evidence that mitonuclear compatibility influences sexual signals and reproductive outcomes. These findings support the idea that female choice can act as a selective filter against mitonuclear incompatibility, linking metabolic coadaptation to the evolution of sexual signalling.

## 2. Materials and Methods

### Drosophila strains and maintenance

A unique panel of 81 *Drosophila melanogaster* mitonuclear genotype combinations were used [24]. This panel was produced through full factorial crossing of 9 isogenic lines, which were acquired from the “Global Diversity Panel” [25]. The isogenic lines in the panel are **A**/ZIM184 (Zimbabwe), **B**/B04 (Beijing), **C**/I16 (Ithaca), **D**/I23 (Ithaca), **E**/N14 (Netherlands), **F**/N15 (Netherlands), **G**/T01 (Tasmania), **H**/T23 (Tasmania), **i**/N01 (Netherlands). The lines have varying phylogeographic relatedness and are representative of the global genetic diversity of fruit fly populations, making results more generalisable. There are nine coadapted and seventy-two disrupted mitonuclear genotypes, allowing systematic investigation into mitonuclear interactions. Each coadapted line has its nuclear and mitochondrial genomes coming from the same geographical population as each other, while disrupted lines pair nuclear and mitochondrial genomes from different populations. We use a 2-letter code to denote all possible mitonuclear genotypes. The first letter corresponds to the nuclear genome, with the second letter referring to the mtDNA. For example, genotype “FC” has the nuclear genotype from strain “F” and the mtDNA from strain “C”.

Throughout this paper we distinguish two related but distinct concepts: **mitonuclear coadaptation** refers to within-individual co-evolution of the nuclear and mitochondrial genomes (ie: whether a fly’s own genomes are coadapted), while **mitonuclear compatibility** refers to between-individual concordance of mitonuclear backgrounds between a mating pair (ie: whether male and female carry matching nuclear and/or mitochondrial backgrounds). These are independent axes of variation in our design: a coadapted male may or may not be compatible with a given female, and an incompatible male may nonetheless carry coadapted genomes. The individual mitonuclear combinations were produced using a balancer chromosome crossing scheme (for sull scheme see Supplementary Figure 1 of reference [24]). Balancer chromosomes are non-recombining and homozygous lethal chromosomes. They can be used to replace the nuclear background that mitochondria interact with when crossed with the isogenic lines. Only 80 genotypes were used for experiments, as the mitonuclear genotype F_nuc_xA_mito_ suffers for a lethal mitonuclear incompatibility and consequently was not viable. Previous sequencing data found no evidence for endosymbionts like *Wolbachia* present in any of the fly genotypes used to make the panel [25].

The flies were maintained on sugar-yeast-agar (SY) food, at 25°C and 50% humidity, on a 12:12 hour light:dark cycle (see Supplementary Table 1 for recipe). Flies from the 80 viable lines were reared in two sets of sequential lays. The first lay was 36 hours long and was initiated using 10-50 parental flies from each line into an unused food vial. Following the initial lay, the flies were placed on new food for 24 hours, and these were used to propagate the next generation and for the experiments. Propagation lays were left for 12 days to allow flies to hatch. Vials were density controlled, having under 120 larvae, and had a standardised amount of live yeast granules.

### Cuticular Hydrocarbon (CHC) Assay

We performed the CHC screen using virgin males, which were collected from each mitonuclear genotype within 8 hours of eclosion and aged in same-genotype vials for 5 days. At this point, three males from each genotype were anaesthetised with CO_2_ and placed in 4 mL glass vials. A thin layer of tissue paper (Kimwipe) was placed on the CO_2_ gas pad and changed for every genotype to avoid cross-contamination of CHCs. CHCs were extracted by washing with 100 µl of hexane (HPLC grade, 99%, Fischer Scientific) containing 100 µM of decane (99+% Acros organics) as an internal standard. Flies were lightly vortexed and incubated for 5 minutes at room temperature. The hexane was removed to a 200 µL glass insert in 2 mL glass vials and capped with PTFE septa caps. Samples were either immediately processed or stored at –80 °C (for no longer than 1 day) until processing; this is due to the sample limit of the GCMS.

Extracts were analysed using a method adapted from Dembeck et al. 2015 [26] using an Agilent Technologies 8890 gas chromatograph (GC) with a 5977B mass spectrometry detector (MSD). 1 µL of sample was introduced using a 10 µL syringe by an automated liquid sampler into a 900 µL splitless liner at 290 °C. The purge flow to the split vent was set to 100 mL/min at 1 min. The initial oven temperature was 50 °C and held for 1 minute before a ramp of 20 °C/min up to 150 °C, a second ramp of 5 °C/min to a final temperature of 300 °C and a final hold of 10 minutes. Separation was performed on a DB-5ms capillary column (40 m x 250 μm x 0.25 μm) with a flow rate of 1.1 mL/min under constant flow. Helium was used as the carrier gas. Mass spectral data were acquired using an extraction ion source. Ion source temperature was 230 °C, quadrupole temperature 150 °C and MSD transfer line was 300 °C. Ions were detected in the 30-600 m/z range, equating to a scan speed of 1.4 scans/s, with a delay time of 5 mins. Data were processed in Agilent Masshunter Qualitative and Quantitative software with compound identification performed by comparison to the Wiley NIST 14 database. These were then manually curated to ensure accurate identification. Chiral compounds and position of double bonds were not investigated and as such peak assignments are not definitive.

Male carcasses were kept for all samples at –80 °C to correct CHC production with body mass. Following the CHC sample collection, all male carcasses were dried in an oven set at 65 °Cfor 16 hours (overnight). Dry weight was obtained for all fly triplets using a Cubis II Satorious microbalance.

### Mating Success & Latency to Mating

This experiment was run over four consecutive generations (“blocks”), with each generation contributing equal number of flies across the panel.

We examined latency to mating, with individual females presented males having either fully compatible, or disrupted mitonuclear backgrounds with respect to the female’s own genotype (**Table 1**). Every generation we collected virgin males and females from the mitonuclear panel. Specifically, we collected virgin females with coadapted mitonuclear genotypes (9 genotypes in total), and male genotypes spanning the whole panel. These flies were kept in sex- and genotype-specific vials and aged for 4 days.

**Table 1:** Experimental setup for mate choice experiment. Here we how all the possible genotypes BB females get exposed to. This setup is consistent for all female genotypes across our 9×9 Drosophila panel.

| Male treatment | Female Genotype<br>(nuDNA-mtDNA) | Possible male genotypes presented<br>(nuDDNA-mtDNA) |
| --- | --- | --- |
| 1. matched | BB | **BB** |
| 2. matching nuDNA | BB | **BA, BC, BD, BE, BF, BG, BH, Bi** |
| 3. matching mtDNA | BB | **BB, CB, DB, EB, FB, GB, HB, iB** |

Mating arenas were built out of tissue culture 24-well plates, with small holes drilled on the lid corresponding to the middle of each well; these fit one individual fly and were covered with a small foam square to prevent escapes. For each experimental run, we started by placing females of a given genotype into each well of the plate using an insect aspirator. Four plates were setup every run of the experiment giving a total of 96 females, with three runs being performed each day (we measured over two consecutive days per block). Once all females were in their chambers, we added a single male to each chamber. Males can be classified in three different categories:

1. Male’s mitonuclear background is fully compatible with the female (male is also coadapted) ie: BB female × BB male
2. Only the nuclear background is compatible with the female (nuclear compatibility only; disrupted mitonuclear genotype). Mitochondrial genotype is sourced randomly covering all possible 9 combinations. ie: BB female × BX male
3. Only the mitochondrial background is compatible with the female (mitochondrial compatibility only; disrupted mitonuclear genotype). Nuclear genotype is sourced randomly covering all possible 9 combinations. ie: BB female × XB male

Given the complexity of the experimental design, we randomised male groups 2 and 3 to cover all possible genotype combinations (see Table 1 for further explanation).

We setup each 24-well plate within 5 minutes and once all males were added, we placed them in a well-lit quiet chamber with two recording cameras (one camara recorded 2 plates). Flies were recorded for 2 hours and then discarded after this recording period. The 24-well plates were then washed with deionized water plus ethanol to clean and remove any pheromones possibly affecting the subsequent runs.

### Statistical Analyses

Raw CHC peak areas were first normalised to the internal standard (100 µM decane) included in each sample. Because CHCs form compositional data (each compound representing a part of a total blend), peak areas were converted to proportions and transformed using the centred log-ratio (CLR) transformation. This approach removes the unit-sum constraint and maps the proportional CHC blend into Euclidean space (dimension n − 1), allowing standard multivariate analyses [27, 28]. Covariate adjustment for block and two-thirds of body mass was then performed by linear regression in CLR space, preserving Aitchison geometry, and the resulting residuals were used for all subsequent analyses. We assessed the significance of nuclear background, mitochondrial haplotype, and their interaction on overall CHC composition using permutational multivariate analysis of variance (PERMANOVA) based on Eucledian distances with 9,999 permutations (R package vegan). To further evaluate homogeneity of multivariate dispersion among genotypes, we used PERMDISP. We also conducted MANOVA with residual randomisation permutation procedures (RRPP; R package geomorph) to test for genotype effects while accounting for within-group covariance. Note that R^2^ values are not directly comparable across methods because PERMANOVA partitions dissimilarity, whereas RRPP MANOVA partitions variance. To visualise differences in CHC blend composition among genotypes, we performed principal components analysis (PCA) on the CLR-transformed data. Scores from the constrained ordination were used to identify individual CHCs contributing most strongly to interaction-structured differences in blend composition.

For mating assays, we analysed two different traits. The first trait was the ability to mate within the 2-hour timeframe, which is binary response variable. To analyse the ability to copulate, we used generalised linear mixed models (GLMM), with mating as a response variable. Mating category and female nuclear genetic background were modelled as fixed effects and “run” nested within experimental block was modelled as a random effect. For males that successfully copulated with the female, we measured latency to mating. Here, we used GLMMs with poisson distribution to model time to copulation as a response variable, with mating category and female nuclear background as fixed effects. Run nested within experimental block was modelled as a random effect. We noticed that data was overdispersed and consequently we included in the models an observation-level random effect.

To test whether mitonuclear-structured variation in CHC blend composition predicts copulation probability, we linked the two datasets at the genotype level. For each of the 80 male genotypes, mean PC scores (PC1–PC5) from the CLR-transformed CHC profiles were computed and joined to the individual-level mating observations. A logistic GLMM was then fitted with copulation success as the binary response, PC1–PC5 as additional fixed-effect predictors alongside female nuclear background, and run nested within week as a random effect, matching the structure of the original mating model. A likelihood ratio test against a null model retaining only female background and the random effect assessed whether CHC axes collectively explained variance in mating outcome. To obtain an out-of-sample estimate of predictive accuracy, we performed leave-one-genotype-out cross-validation (LOGO-CV). More specifically, for each of the 80 genotypes in turn the model was fitted on the remaining 79 genotypes and used to predict copulation probability for the held-out genotype (population-level prediction, marginalising over random effects). Pearson *r* and Spearman ρ between predicted and observed mating rates across genotypes were used to quantify out-of-sample predictive accuracy. Because CHC profiles were measured from pooled triplets, all individuals of the same male genotype share identical PC predictor values; the LOGO-CV accounts for this by holding out entire genotypes rather than individual observations. All analyses were performed in R using the *lme4* package. To test whether CHC effects on mating are compatibility-dependent rather than absolute, we extended the model by adding mating category × PC interaction terms (Type × PC1–PC5), using mating category (Match, NucMatch, MitoMatch) as a proxy for between-individual mitonuclear compatibility. A likelihood ratio test compared this interaction model against the additive model. To further assess whether predictive accuracy varies with compatibility context, we repeated the LOGO-CV separately within each mating category stratum; the co-evolved (Match) stratum was excluded from this analysis because only nine co-evolved genotypes are present, leaving training sets too small for reliable model fitting after holding one out.

## 3. Results

### Cuticular hydrocarbons

We identified 21 cuticular hydrocarbons across the 80 mitonuclear genotypes screened (see Supplementary table 2 for the list of compounds). We analysed CHC data by looking at the compositional blend (obtaining the log-ratio of the compounds) per mitonuclear genotype.

Distance-based partitioning attributed most explainable multivariate variation in CHCs to nuclear background (PERMANOVA R^2^ = 0.57) with a smaller mitochondrial genome effect (R^2^ = 0.02) and a non-additive interaction component (R^2^ = 0.088 of the total; all p ≤ 0.001). PCA of centred residuals visualised this structure (**Figure 1, Figure 2**): samples clustered by nuclear genotype with mitochondrial haplotypes shifting those clusters as expected for an interaction. An RRPP MANOVA detected robust location shifts for nuclear (R^2^ = 0.091, F = 16.78, p < 0.001) mitochondrial (R^2^ = 0.021, F = 3.84, p < 0.001), and nuclear × mitochondrial (R^2^ = 0.088, F = 2.07, p < 0.001), indicating genuine differences in mean blend composition across genotypes. Multivariate dispersion was homogeneous across groups in CLR space, supporting a location rather than spread difference. To identify CHCs contributing most to the interaction in CLR space, we examined species scores from the interaction-constrained ordination. The largest contributors were Component 19 (heptacosene) and Component 16 (pentacosene), highlighting these compounds as key drivers of mitonuclear-structured differences in the blend.

**Figure 1.**
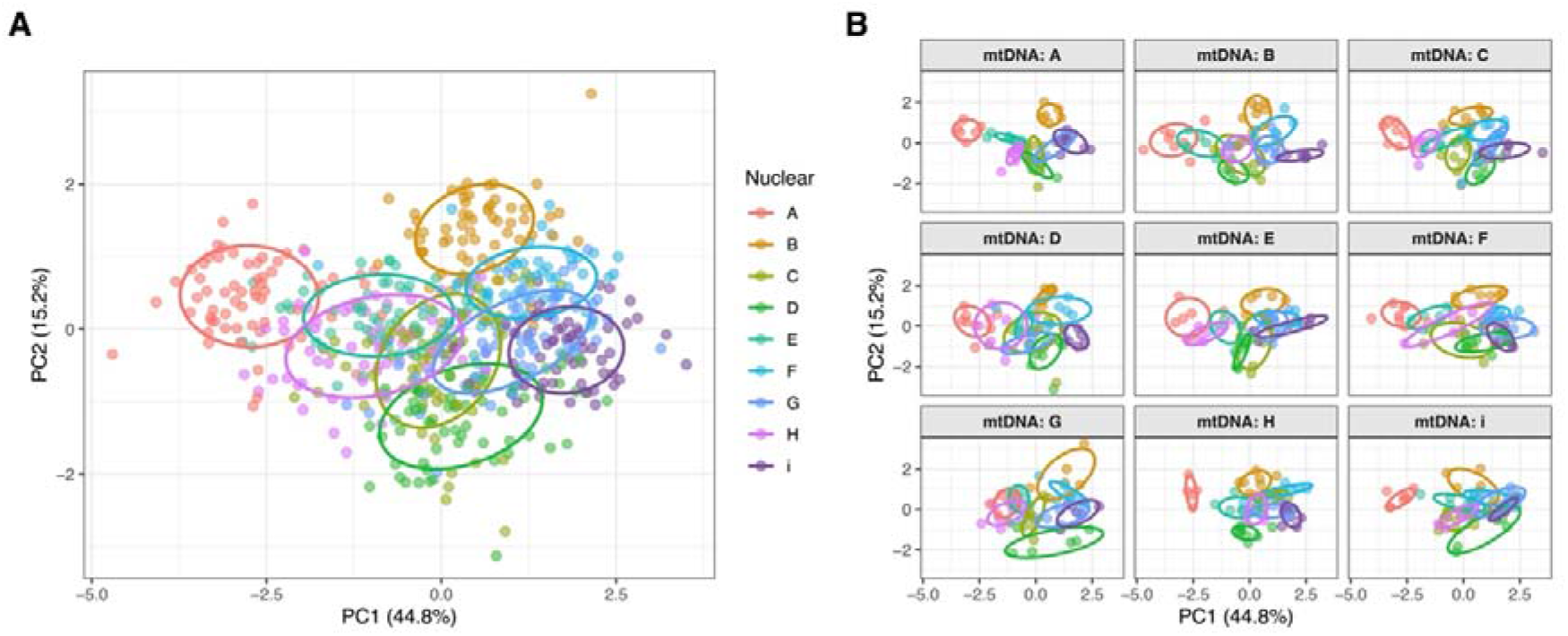
Principal component analysis (PCA) of cuticular hydrocarbon (CHC) profiles. **(A)** PCA plot coloured by nuclear background, with ellipses indicating the ± 1SD of each genotype. This panel illustrates the overall separation of nuclear genomes in multivariate CHC space. **(B)** The same PCA scores, but faceted by mitochondrial haplotype (mtDNA genome), highlighting how nuclear genotypes cluster within each mitochondrial background. Axis labels indicate the proportion of variance explained by PC1 and PC2.

**Figure 2:**
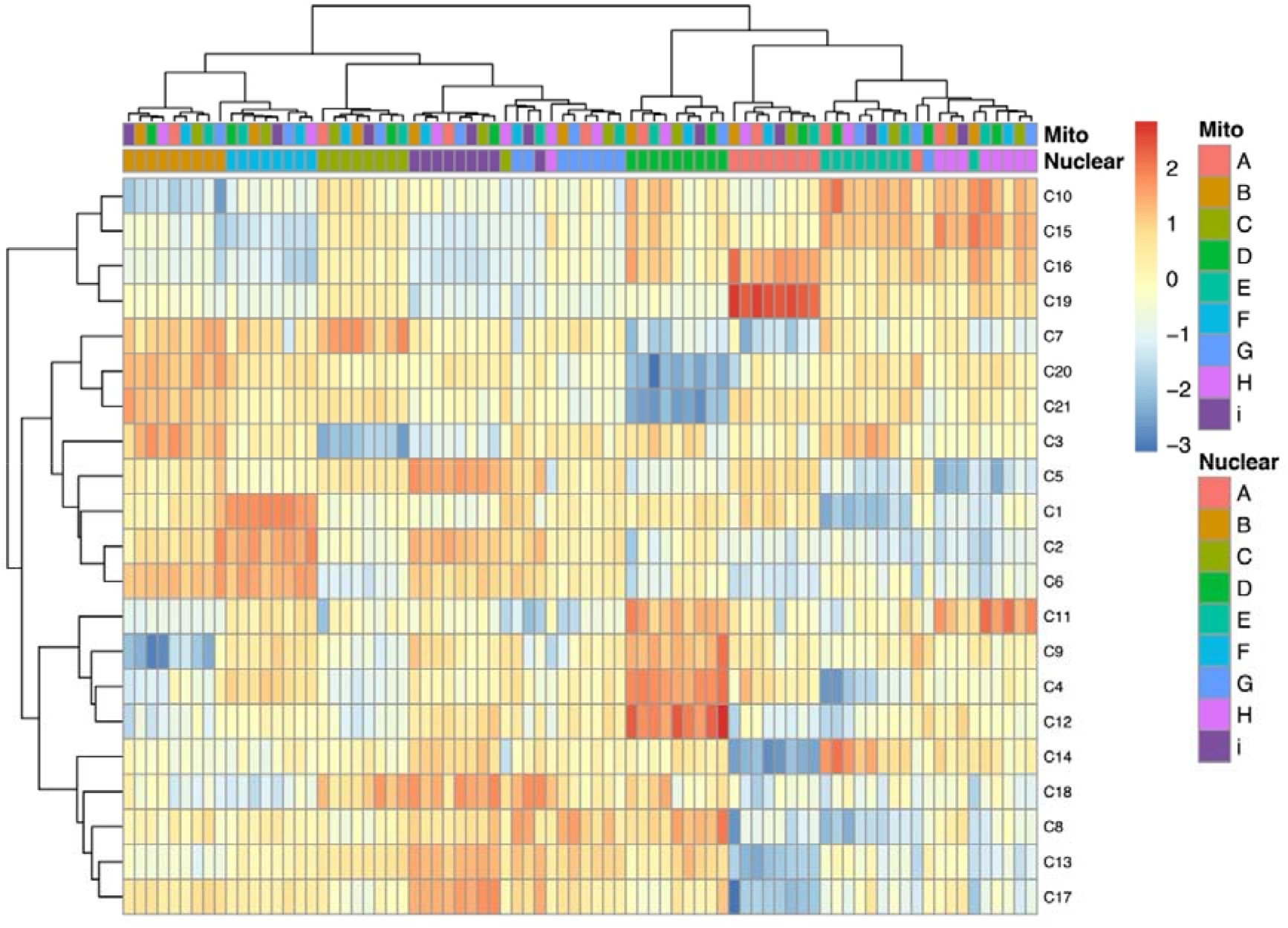
Heatmap of mean cuticular hydrocarbon (CHC) profiles across mitonuclear genotypes. Rows correspond to individual CHC compounds and columns represent the mean profile of each mitonuclear combination after adjustment for body mass and block effects. Columns are annotated by nuclear background and mitochondrial haplotype. Hierarchical clustering of compounds (Ward’s method) highlights patterns of co-variation among hydrocarbons, while column ordering reflects nuclear and mitochondrial grouping. For full list of CHC Identifiers see Supplementary Table 2.

### Mating success & latency to mating

We find an overall effect of male genotype category (χ^2^ = 22.99, p < 0.001), and an effect of female genotype (χ^2^ = 49.29, p < 0.001). No significant interactive effect was observed in our models; hence we did not include this interaction in our models. We then extracted the marginal means from the binomial models to examine the how the three mating categories influenced copulation success. We find that fully compatible males (coadapted genomes paired with a genotypically identical female) had higher mating success (0.50 [0.38,0.63]) compared to those with only nuclear compatibility (0.37 [0.26, 0.49]) or only mitochondrial compatibility (0.31 [0.21, 0.43], **Figure 3A**).

**Figure 3:**
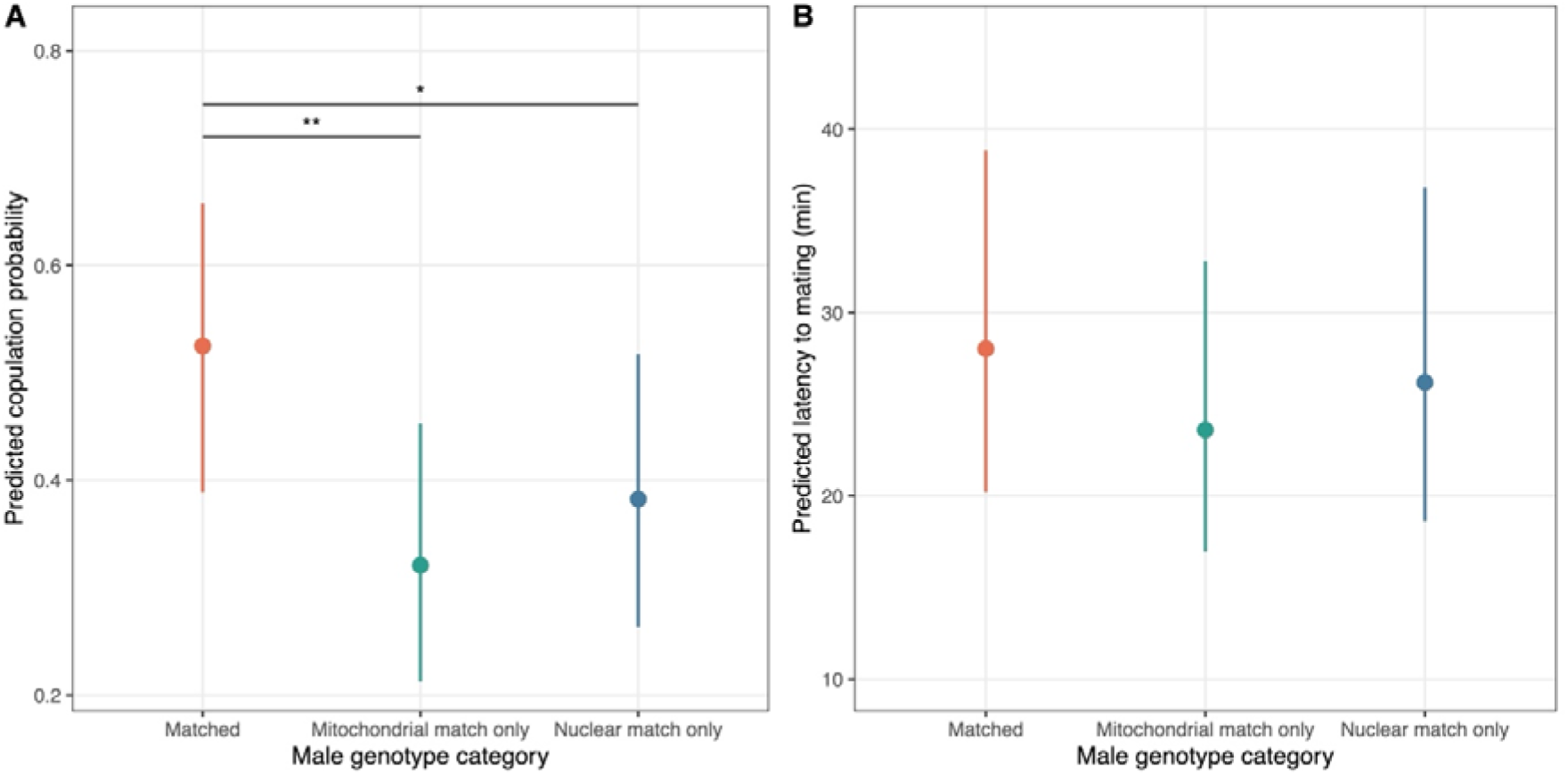
Marginal means (± 95% CI) output from mixed effects models. **(A)** Results from mating success analysis where we examine the number of successful copulations within a 2-hour period across our male genotype treatments. **(B)** Latency to mating (minutes) for pairs that were successful in copulation, across all male genotype treatments.

Our second experiment examined in the group that successfully mated, if there was a difference in latency to mating. Our analysis shows that there is a significant female genotype effect (χ^2^ = 33.70, p < 0.001), however we did not detect a significant male genotypic category (χ^2^ = 3.88, p = 0.1437), nor an interaction between both factors (**Figure 3B**).

### CHC-mediated prediction of compatibility

To directly test whether these mating differences are linked to CHC blend composition, we modelled genotype-level CHC PC scores as predictors of copulation probability using a cross-validated logistic GLMM. CHC axes significantly predicted mating outcome (likelihood ratio test: χ^2^ = 37.66, df = 5, p < 0.001), with PC3 and PC4 emerging as the strongest individual predictors (both p < 0.001, **Figure 4A**). Leave-one-genotype-out cross-validation confirmed that this predictive relationship holds out-of-sample (Pearson r = 0.43; **Figure 4B**), supporting a quantitative link between mitonuclear-structured CHC blend composition and copulation probability. Critically, a model incorporating mating category × CHC interaction terms fit significantly better than the additive model (likelihood ratio test: χ^2^= 40.63, df = 12, p < 0.001) demonstrating that CHC effects on copulation success are compatibility-dependent rather than absolute. The strongest interactions involved the mito-matched category: MitoMatch×PC4 (p = 0.023) and MitoMatch×PC5 (p = 0.049), indicating that these CHC axes predict mating differently when males and females share mitochondrial but not nuclear backgrounds, relative to co-evolved pairs. Nuclear-match interactions were non-significant across all CHC axes. Stratified LOGO-CV was evaluable for the two mismatched categories only (the nine co-evolved genotypes are too few to support leave-one-out validation after holding one out) and showed that CHC blend composition predicted mating more accurately in mito-matched pairs (Pearson r□=□0.48) than nuclear-matched pairs (r□=□0.34), consistent with mtDNA-structured variation in the CHC blend carrying particular information about compatibility in that context. Together, these results indicate that CHC blend composition reflects mitonuclear compatibility in a way that is detected by females, and that the relevant information is context-dependent rather than a fixed quality signal.

**Figure 4:**
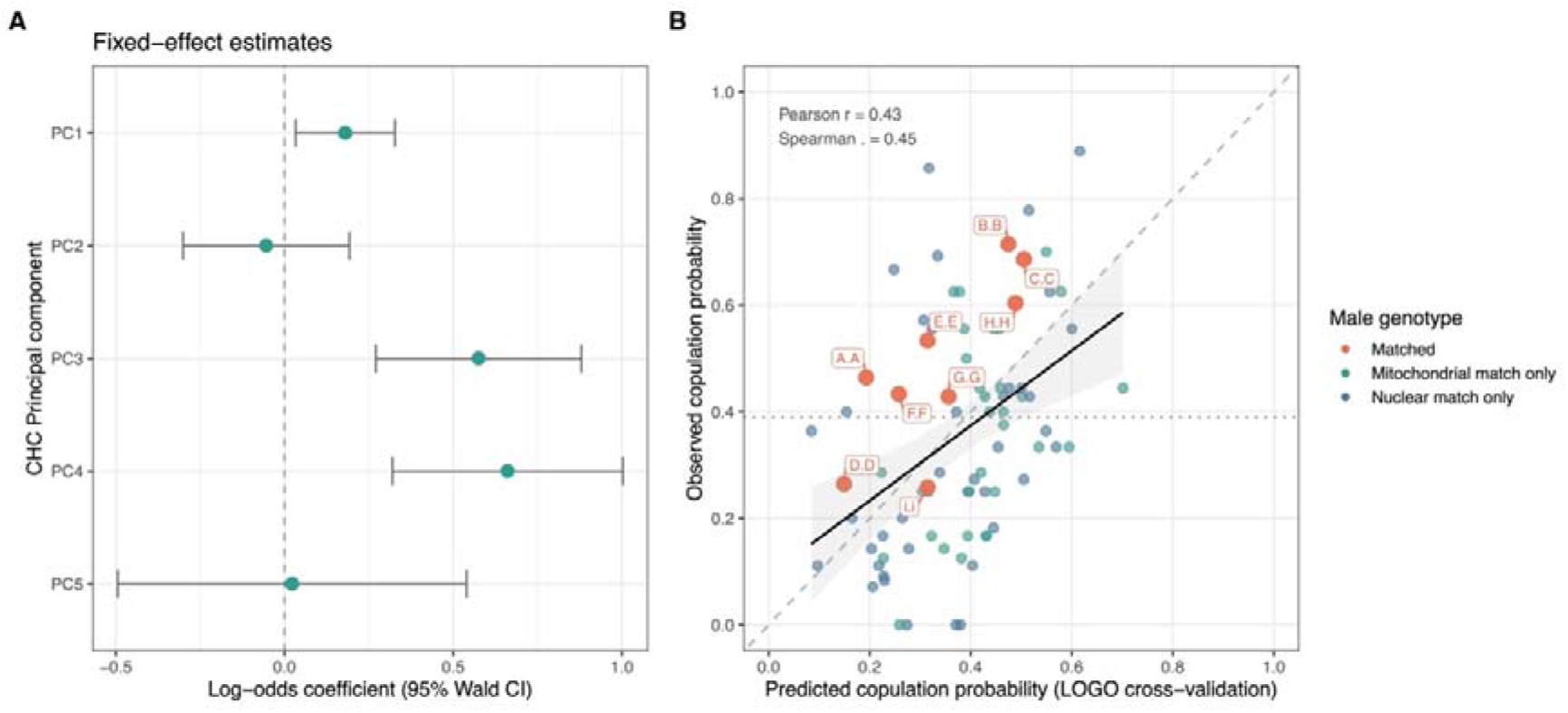
CHC blend composition predicts copulation probability across the mitonuclear panel. **(A)** Fixed-effect log-odds coefficients (± 95% Wald CI) from a logistic GLMM predicting copulation probability from genotype-mean PC scores of CLR-transformed CHC profiles, with female identity and run-within-week as covariates. PC axes with confidence intervals excluding zero (PC1, PC3, PC4) indicate CHC dimensions that significantly predict mating success. **(B)** Cross-validated prediction of copulation probability. Each point represents one of the 80 male genotypes; the x-axis shows predicted copulation probability from leave-one-genotype-out cross-validated model fits, and the y-axis shows observed copulation rate in no-choice assays. The dashed line indicates perfect prediction; the dotted horizontal line marks the overall mean copulation rate. The solid line shows the linear trend with 95% confidence interval. Pearson r and Spearman ρ reflect out-of-sample predictive accuracy across all 80 genotypes.

## 4. Discussion

Our results show that nuclear background, mtDNA haplotype, and their interaction modulate the blend of cuticular hydrocarbons (CHCs) in *Drosophila* males. Behaviourally, these chemical differences do matter: males carrying coadapted mitonuclear genomes achieve higher copulation success when paired with a genotypically compatible female, possibly linking genetic architecture to a composite chemical signal with downstream consequences on reproductive outcomes. This pattern aligns with the view that CHCs are heritable, polygenic traits whose components covary due to shared biosynthetic steps with co-regulated upstream resource supply (acetyl-CoA, ATP, NADPH, etc).

Considering the wider literature, the genotype-CHC-mating relationships we document are not limited to *Drosophila melanogaster*. Across several fruit fly species, abundant standing genetic variation has been repeatedly documented in multivariate CHC space [12, 26, 29, 30]. This variation has clear functional consequences, as manipulation of single genes (for example desaturases), can alter both the production and perception of pheromonal blends, highlighting a close genetic coupling between signal and receiver [31]. Patterns observed between natural populations reinforce this link, as divergence in female CHC profiles often coincides with assortative mating and the early stages of reproductive isolation [32–34]. Direct causal evidence comes from perfuming experiments, in which the transfer of donor hydrocarbons predictably modifies mating success, demonstrating that CHC blends themselves (rather than correlated morphological or behavioural traits) mediate mate choice [32].

Comparable dynamics are evident across a wide range of other taxa. In crickets, genetically distinct inbred lines differ in CHC composition, genotypes reshape chemical profiles, and field-based estimates reveal both directional and stabilising sexual selection acting on multivariate CHC axes [35]. In parasitoid wasps, CHC expression reflects joint effects of wasp genotype and host environment, with measurable behavioural consequences such as altered aggression by ants [36, 37]. In social insects, colony- and family-specific CHC signatures underpin nestmate recognition, dominance interactions and reproductive differentiation [38, 39]. Even outside insects, spiders exhibit family-specific cuticular profiles that function as cues for kin recognition or social interaction [40, 41]. Collectively, these findings point to a general principle: genetic variation generates structured diversity in CHC blends, and selection operates on that structure through its effects on social and reproductive behaviour.

Our mitonuclear perspective offers a simple explanation for why genetic effects on CHCs align so closely with mating outcomes. Cuticular hydrocarbon biosynthesis is metabolically demanding [23]: it depends on acetyl-CoA supply, ATP, and NADPH, and is carried out by nuclear-encoded enzymes whose effective activity depends on mitochondrial performance [42, 43]. If efficient fatty-acid elongation and desaturation require coordinated function between nuclear and mitochondrial genomes, then disrupted mitonuclear combinations within a male are expected to alter the relative composition CHC blends and possibly reducing overall signal output. This constraint is especially clear in oenocytes, the specialised cells responsible for CHC synthesis. Disrupting fatty-acid metabolism in these cells compromises cuticular waterproofing and can be lethal [44, 45], highlighting both the cost and tight metabolic integration of CHC production. Because CHCs also play a central role in mate recognition [46, 47], variation in blend composition provides females with information about male metabolic coordination. Under this view, female preference for males with coadapted mitonuclear genomes - and whose backgrounds are compatible with the female’s own - arises naturally, as CHC blends act as honest indicators of how well nuclear and mitochondrial genomes function together [48].

The two compounds contributing most strongly to the mitonuclear interaction axis are 7-heptacosene (Compound 19, C27:1) and 7-pentacosene (Compound 16, C25:1). Both are male-specific monoenes whose relative proportions show well-documented genotype and environment dependence in the wider literature [49, 50]. For example, 7-pentacosene (7-P) is one of the two principal male pheromones in *D. melanogaster*, alongside 7-tricosene, and its relative abundance varies systematically with geoclimatic parameters (ie: latitude, longitude, mean temperature and vapour pressure), suggesting it is sensitive to metabolic state and environmental resource availability [49]. A dedicated QTL region, *sept*, has been mapped specifically to variation in male 7-P proportions, and temperature selection experiments demonstrate that the 7-P ratio is phenotypically plastic and co-varies with desiccation resistance [26]. For 7-heptacosene, population-level variation has been documented across *D. yakuba* populations from different African islands, where it co-varies with mating rate, and its biosynthesis is directly downstream of the elongase *eloF*; a nuclear-encoded enzyme responsible for producing C27 and C29 compounds whose knockdown shifts the blend towards shorter C23 and C25 hydrocarbons [51]. The fact that both compounds sit at the end of the fatty-acid elongation chain and are therefore most sensitive to the efficiency of upstream biosynthetic steps, makes them plausible candidates for reflecting mitonuclear metabolic coordination. Their emergence as the strongest drivers of the interaction axis is thus mechanistically coherent: if disrupted mitonuclear males have compromised elongase and desaturase efficiency, it is the longer-chain, later-synthesised compounds that would be expected to shift most.

Cuticular hydrocarbons serve dual functions in insects, contributing both to sexual communication and to water balance [10]. As a result, selection acting on physiology can directly affect the sexual signal. For example, shifts towards longer or more saturated hydrocarbons can improve desiccation resistance [52], but at the same time move individuals to a different position in chemical signal space as perceived by choosy mates [11]. Environmental stressors that limit energy availability or disrupt metabolic balance (such as temperature extremes or dietary imbalance) are therefore expected to accentuate genetic effects on CHC composition and increase mating differences among genotypes [53, 54]. Consistent with this expectation, nutritional geometry experiments in field crickets show that macronutrient intake jointly shapes CHC expression, desiccation resistance, and male mating success, revealing partial alignment between natural and sexual selection acting on CHCs [55]. Together, these patterns indicate that CHC expression is inherently condition-dependent, integrating genetic, physiological, and environmental variation into signals that are simultaneously shaped by ecology and mate choice.

These connections point to clear next steps that would sharpen inference and deepen our understanding of the mechanisms involved. Causality could be tested by directly manipulating the compounds that load most strongly on the mitonuclear interaction axis, for example through synthetic add-backs to males with disrupted mitonuclear genomes, and asking whether compatibility-dependent mating success is restored. Mechanistic insight could then be gained by linking variation in CHC blends to underlying metabolic processes, such as oenocyte expression of elongases and desaturases or measures of mitochondrial performance and redox balance, and testing whether these biochemical traits co-vary with the CHC axes that differentiate genotypes. The robustness of these relationships could be assessed by introducing environmental challenges, such as dietary or thermal stress, to test predicted genotype-by-environment amplification of both CHC variation and mating outcomes. Finally, multivariate and causal analytical approaches that treat CHC axes as phenotypes rather than individual compounds could help identify which aspects of blend structure most reliably predict mate choice. Furthermore, we only selected one timepoint for our CHC measurements, whereas we know CHC production is a dynamic process, especially during courtship [56]. If mate choice relies on such context-dependent cues, pooled-body profiles may dilute the most behaviourally relevant signal. Future work using time-resolved sampling or fly-specific extractions would therefore help refine the mapping from genotype to the components of the CHC mixture that are most directly involved in mate choice.

In sum, by pairing multivariate analysis with behavioural assays, we show that the mitonuclear genotype, structures the chemical signals that females use to choose mates. This places a conspicuous sexual signal within a well-established framework of cellular bioenergetics and genetic compatibility, offering a simple mechanism by which sexual selection can help maintain co-adapted mitonuclear combinations and shape patterns of adaptation across environments. More broadly, it suggests that cuticular hydrocarbons are best understood as integrated, metabolically grounded traits whose genetic architecture, ecological function, and behavioural consequences are closely linked.

## Acknowledgements

MFC is funded by a Natural Environment Research Council Fellowship (NE/V014307/1) and a Leverhulme Trust grant (RPG-2023-198).

## Author Contributions

TA, NL, & MFC conceived the ideas. TA & MFC designed the methodology and analyzed the data. TA & SH collected and analyzed GCMS data. TA collected all mating assay data. MFC led the writing of the manuscript. All authors contributed critically to the drafts and gave final approval for publication.

## Data Accessibility Statement

All data will be made available following publication. All R scripts and R-markdown documents are available with this manuscript submission.

## References

1. Darwin, C., The descent of man, and selection in relation to sex, in The descent of man, and selection in relation to sex. 2008, Princeton University Press.

2. Zahavi, A., Mate selection-a selection for a handicap. J Theor Biol, 1975. 53(1): p. 205–14.

3. Andersson, M., Sexual selection, in Sexual Selection. 2019, Princeton University Press.

4. Grafen, A., Biological signals as handicaps. Journal of Theoretical Biology, 1990. 144(4): p. 517–546.

5. Cotton, S., K. Fowler, and A. Pomiankowski, Do sexual ornaments demonstrate heightened condition-dependent expression as predicted by the handicap hypothesis? Proceedings of the Royal Society of London. Series B: Biological Sciences, 2004. 271(1541): p. 771–783.

6. Hamilton, W.D. and M. Zuk, Heritable true fitness and bright birds: a role for parasites? Science, 1982. 218(4570): p. 384–7.

7. Folstad, I. and A.J. Karter, Parasites, Bright Males, and the Immunocompetence Handicap. The American Naturalist, 1992. 139(3): p. 603–622.

8. Martínez-Lendech, N., et al., Sexual signals reveal males’ oxidative stress defences: Testing this hypothesis in an invertebrate. Functional Ecology, 2018. 32(4): p. 937–947.

9. Moore, F.R., D.M. Shuker, and L. Dougherty, Stress and sexual signaling: a systematic review and meta-analysis. Behavioral Ecology, 2015. 27(2): p. 363–371.

10. Howard, R.W. and G.J. Blomquist, Ecological, behavioral, and biochemical aspects of insect hydrocarbons. Annu Rev Entomol, 2005. 50: p. 371–93.

11. Blows, M.W., Interaction between natural and sexual selection during the evolution of mate recognition. Proceedings of the Royal Society of London. Series B: Biological Sciences, 2002. 269(1496): p. 1113–1118.

12. Foley, B., et al., Natural genetic variation in cuticular hydrocarbon expression in male and female Drosophila melanogaster. Genetics, 2007. 175(3): p. 1465–77.

13. Thomas, M.L. and L.W. Simmons, Male-derived cuticular hydrocarbons signal sperm competition intensity and affect ejaculate expenditure in crickets. Proc Biol Sci, 2009. 276(1655): p. 383–8.

14. Bontonou, G. and C. Wicker-Thomas, Sexual Communication in the Drosophila Genus. Insects, 2014. 5(2): p. 439–58.

15. Lane, S.M., et al., Sexual Selection on male cuticular hydrocarbons via male-male competition and female choice. J Evol Biol, 2016. 29(7): p. 1346–55.

16. Holze, H., L. Schrader, and J. Buellesbach, Advances in deciphering the genetic basis of insect cuticular hydrocarbon biosynthesis and variation. Heredity (Edinb), 2021. 126(2): p. 219–234.

17. Mitchell, P., Chemiosmotic coupling in oxidative and photosynthetic phosphorylation. Biological Reviews, 1966. 41(3): p. 445–501.

18. Barrell, B.G., A.T. Bankier, and J. Drouin, A different genetic code in human mitochondria. Nature, 1979. 282(5735): p. 189–94.

19. Burton, R.S. and F.S. Barreto, A disproportionate role for mtDNA in Dobzhansky-Muller incompatibilities? Molecular Ecology, 2012. 21(20): p. 4942–57.

20. Rand, D.M., R.A. Haney, and A.J. Fry, Cytonuclear coevolution: the genomics of cooperation. Trends Ecol Evol, 2004. 19(12): p. 645–53.

21. Hill, G.E., et al., Plumage redness signals mitochondrial function in the house finch. Proc Biol Sci, 2019. 286(1911): p. 20191354.

22. Hill, G.E. and J.D. Johnson, The mitonuclear compatibility hypothesis of sexual selection. Proceedings of the Royal Society B: Biological Sciences, 2013. 280(1768): p. 20131314.

23. Berson, J.D. and L.W. Simmons, A costly chemical trait: phenotypic condition dependence of cuticular hydrocarbons in a dung beetle. J Evol Biol, 2018. 31(12): p. 1772–1781.

24. Carnegie, L., et al., Mother’s curse is pervasive across a large mitonuclear Drosophila panel. Evolution Letters, 2021. 5(3): p. 230–239.

25. Grenier, J.K., et al., Global Diversity Lines–A Five-Continent Reference Panel of Sequenced Drosophila melanogaster Strains. G3 Genes|Genomes|Genetics, 2015. 5(4): p. 593–603.

26. Dembeck, L.M., et al., Genetic architecture of natural variation in cuticular hydrocarbon composition in Drosophila melanogaster. Elife, 2015. 4.

27. Hine, E., S.F. Chenoweth, and M.W. Blows, Multivariate quantitative genetics and the lek paradox: genetic variance in male sexually selected traits of Drosophila serrata under field conditions. Evolution, 2004. 58(12): p. 2754–62.

28. Sztepanacz, J.L. and H.D. Rundle, Reduced genetic variance among high fitness individuals: inferring stabilizing selection on male sexual displays in Drosophila serrata. Evolution, 2012. 66(10): p. 3101–10.

29. Oliveira, C.C.d., et al., Variations on a theme: diversification of cuticular hydrocarbons in a clade of cactophilic Drosophila. BMC Evolutionary Biology, 2011. 11(1): p. 179.

30. Sharma, M.D., et al., The Genetics of Cuticular Hydrocarbon Profiles in the Fruit Fly Drosophila simulans. Journal of Heredity, 2012. 103(2): p. 230–239.

31. Marcillac, F., Y. Grosjean, and J.F. Ferveur, A single mutation alters production and discrimination of Drosophila sex pheromones. Proc Biol Sci, 2005. 272(1560): p. 303–9.

32. Davis, J.S., et al., A shift to shorter cuticular hydrocarbons accompanies sexual isolation among Drosophila americana group populations. Evol Lett, 2021. 5(5): p. 521–540.

33. Frentiu, F.D. and S.F. Chenoweth, Clines in cuticular hydrocarbons in two Drosophila species with independent population histories. Evolution, 2010. 64(6): p. 1784–94.

34. Mas, F. and J.-M. Jallon, Sexual Isolation and Cuticular Hydrocarbon Differences between Drosophila santomea and Drosophila yakuba. Journal of Chemical Ecology, 2005. 31(11): p. 2747–2752.

35. Weddle, C.B., et al., Sex-specific genotype-by-environment interactions for cuticular hydrocarbon expression in decorated crickets, Gryllodes sigillatus: implications for the evolution of signal reliability. J Evol Biol, 2012. 25(10): p. 2112–2125.

36. Pokorny, T. and J. Ruther, Cuticular Hydrocarbon Polymorphism in a Parasitoid Wasp. J Chem Ecol, 2023. 49(1-2): p. 36–45.

37. Buellesbach, J., S.G. Vetter, and T. Schmitt, Differences in the reliance on cuticular hydrocarbons as sexual signaling and species discrimination cues in parasitoid wasps. Front Zool, 2018. 15: p. 22.

38. Johnson, B.R., E. van Wilgenburg, and N.D. Tsutsui, Nestmate recognition in social insects: overcoming physiological constraints with collective decision making. Behav Ecol Sociobiol, 2011. 65(5): p. 935–944.

39. Esponda, F. and D.M. Gordon, Distributed nestmate recognition in ants. Proc Biol Sci, 2015. 282(1806): p. 20142838.

40. Weiss, K. and J.M. Schneider, Family-specific chemical profiles provide potential kin recognition cues in the sexually cannibalistic spider Argiope bruennichi. Biol Lett, 2021. 17(8): p. 20210260.

41. Grinsted, L., T. Bilde, and P. d’Ettorre, Cuticular hydrocarbons as potential kin recognition cues in a subsocial spider. Behavioral Ecology, 2011. 22(6): p. 1187–1194.

42. Blomquist, G.J. and M.D. Ginzel, Chemical Ecology, Biochemistry, and Molecular Biology of Insect Hydrocarbons. Annu Rev Entomol, 2021. 66: p. 45–60.

43. Yan, H. and J. Liebig, Genetic basis of chemical communication in eusocial insects. Genes Dev, 2021. 35(7-8): p. 470–482.

44. Wicker-Thomas, C., et al., Flexible origin of hydrocarbon/pheromone precursors in Drosophila melanogaster. J Lipid Res, 2015. 56(11): p. 2094–101.

45. Chiang, Y.N., et al., Steroid Hormone Signaling Is Essential for Pheromone Production and Oenocyte Survival. PLOS Genetics, 2016. 12(6): p. e1006126.

46. Billeter, J.C., et al., Specialized cells tag sexual and species identity in Drosophila melanogaster. Nature, 2009. 461(7266): p. 987–91.

47. Ferveur, J.F., Cuticular hydrocarbons: their evolution and roles in Drosophila pheromonal communication. Behav Genet, 2005. 35(3): p. 279–95.

48. Hill, G.E., Mitonuclear Mate Choice: A Missing Component of Sexual Selection Theory? Bioessays, 2018. 40(3).

49. Rouault, J., P. Capy, and J.M. Jallon, Variations of male cuticular hydrocarbons with geoclimatic variables: an adaptative mechanism in Drosophila melanogaster? Genetica, 2000. 110(2): p. 117–30.

50. Ferveur, J.F. and J.M. Jallon, Genetic control of male cuticular hydrocarbons in Drosophila melanogaster. Genet Res, 1996. 67(3): p. 211–8.

51. Denis, B., A.L. Rouzic, and C. Wicker-Thomas, Hydrocarbon Patterns and Mating Behaviour in Populations of Drosophila yakuba. Insects, 2015. 6(4): p. 897–911.

52. Gibbs, A. and J.G. Pomonis, Physical properties of insect cuticular hydrocarbons: The effects of chain length, methyl-branching and unsaturation. Comparative Biochemistry and Physiology Part B: Biochemistry and Molecular Biology, 1995. 112(2): p. 243–249.

53. Otte, T., M. Hilker, and S. Geiselhardt, Phenotypic Plasticity of Cuticular Hydrocarbon Profiles in Insects. J Chem Ecol, 2018. 44(3): p. 235–247.

54. Baleba, S.B.S., N.-J. Jiang, and B.S. Hansson, Temperature-mediated dynamics: Unravelling the impact of temperature on cuticular hydrocarbon profiles, mating behaviour, and life history traits in three Drosophila species. Heliyon, 2024. 10(17): p. e36671.

55. Simmons, L.W., et al., Nutritional geometry provides insight into the dual roles of natural and sexual selection in insect cuticular hydrocarbon evolution. Functional Ecology, 2025. 39(12): p. 3646–3658.

56. Yew, J.Y., et al., A new male sex pheromone and novel cuticular cues for chemical communication in Drosophila. Curr Biol, 2009. 19(15): p. 1245–54.

