## Supplementary for "Mitonuclear interactions shape male cuticular hydrocarbon profiles with consequences on mating success"

**Table S1:** The *Drosophila* food recipe to make 2 litres of *Drosophila* rearing medium. Nipagin and Propionic acid are included as preservatives.

| **Ingredients** | **ST** |
| --- | --- |
| Agar | 30g |
| Water | 2L |
| Sucrose | 200g |
| Yeast | 200g |
| Nipagin | 60mL |
| Propionic acid | 6mL |

**Table S2: CHC Compound Key.** For manuscript clarity, all CHC compounds identified in our analyses were given a unique identifier.

| **ID** | **Compound** | **Abbrev ID** | **Saturated** | **Branched** |
| --- | --- | --- | --- | --- |
| standard | Decane Results | 10 |  |  |
| C1 | Heneicosane Results | 21:0 | Y | N |
| C2 | 1-Docosene Results | 22:1 | N | N |
| C3 | Z-13-Octadecen-1-yl acetate Results | 21:1:Ester | N | N |
| C4 | Docosane Results | 22:0 | Y | N |
| C5 | 9-Tricosene, (Z)- (1) Results | 23:1 (1) | N | N |
| C6 | 9-Tricosene, (Z)- (2) Results | 23:1 (2) | N | N |
| C7 | 9-Tricosene, (Z)- (3) Results | 23:1 (3) | N | N |
| C8 | Tricosane Results | 23:0 | Y | N |
| C9 | Tetracos-X-ene (1, M) Results | 24:1 (1) | N | N |
| C10 | Tetracos-X-ene (2, M) Results | 24:1 (2) | N | N |
| C11 | Tetracose-X-ene (3, M) Results | 24:1 (3) | N | N |
| C12 | Tetracosane Results | 24:0 | Y | N |
| C13 | 2-Methyltetracosane Results | 25:0:Branched | Y | Y |
| C14 | Z-12-Pentacosene Results | 25:1 (1) | N | N |
| C15 | Z-12-Pentacosene (2, M) Results | 25:1 (2) | N | N |
| C16 | Z-12-Pentacosene (3, M) Results | 25:1 (3) | N | N |
| C17 | Pentacosane Results | 25:0 | Y | N |
| C18 | 2-Methylhexacosane Results | 27:0:Branched | Y | Y |
| C19 | Heptacos-X-ene (M) Results | 27:1 | N | N |
| C20 | Heptacosane Results | 27:0 | Y | N |
| C21 | Octacosane, 2-methyl- Results | 29:0:Branched | Y | Y |
