## Supplementary Code 1 for "Mitonuclear interactions shape male cuticular hydrocarbon profiles with consequences on mating success"

##### Supplementary Pipeline 1: CHC profiling and multivariate analysis

###### Tom Allison, Stuart Harrison, Nick Lane, M. Florencia Camus

#### 2026-03-12

- 1 Overview
- 2 Packages
- 3 Load data
- 4 Internal standard
  correction
- 5 Compositional log-ratio
  transformation and covariate adjustment
- 6 Align metadata
- 7
  Shared colour palette
- 8 Principal component
  analysis
- 9 PERMANOVA
- 10 Dispersion diagnostic
- 11 RRPP MANOVA
- 12 Which CHCs drive the mitonuclear
  interaction?
- 13 Focal compound plots: C19
  (heptacosene) and C16 (pentacosene)
- 14 All-compound interaction
  profiles
- 15 Heatmap of mean CHC
  profiles
- 16 Export manuscript figures to
  PDF
- 17 Session info

### 1 Overview

This pipeline processes raw GC-MS cuticular hydrocarbon (CHC) data
from the *Drosophila melanogaster* mitonuclear panel. It performs
internal standard correction, compositional log-ratio transformation,
covariate adjustment in CLR space, and multivariate analyses (PCA,
PERMANOVA, RRPP) to test for effects of nuclear background,
mitochondrial haplotype, and their interaction on male CHC blend
composition. The analysis also identifies which individual compounds
contribute most strongly to the mitonuclear interaction axis.

---

### 2 Packages

```
library(ggplot2)
library(scales)
library(dplyr)
library(tidyr)
library(vegan)
library(pheatmap)
library(ggrepel)
library(grid)
library(Rmisc)
library(geomorph)
library(patchwork)
library(zCompositions)  # for zero replacement
library(compositions)   # for CLR/ILR
```

---

### 3 Load data

```
dat_path <- params$data_path
CHC.pre  <- read.csv(dat_path, header = TRUE, strip.white = TRUE,
                     stringsAsFactors = FALSE)
CHC.pre  <- CHC.pre[complete.cases(CHC.pre), ]

# Factors
CHC.pre$Nuclear <- factor(CHC.pre$Nuclear)
CHC.pre$Mito    <- factor(CHC.pre$Mito)
CHC.pre$block   <- factor(CHC.pre$block)
```

---

### 4 Internal standard correction

Raw peak areas are normalised to the decane internal standard (column
12) to correct for variation in injection volume and instrument
sensitivity.

```
CHC.cor <- CHC.pre[, 13:33] / CHC.pre[, 12]
CHC     <- cbind(CHC.pre[, 1:10], CHC.cor)
str(CHC)
```

```
## 'data.frame':    555 obs. of  31 variables:
##  $ block         : Factor w/ 21 levels "1","2","3","4",..: 1 1 1 1 1 1 1 1 1 1 ...
##  $ block2        : int  1 1 1 1 1 1 1 1 1 1 ...
##  $ Sample        : int  1 2 3 4 5 6 7 8 9 10 ...
##  $ Nuclear       : Factor w/ 9 levels "A","B","C","D",..: 5 1 3 9 1 2 2 6 5 9 ...
##  $ Mito          : Factor w/ 9 levels "A","B","C","D",..: 1 3 2 9 9 5 6 3 4 1 ...
##  $ Mass          : num  0.582 0.415 0.632 0.512 0.463 0.539 0.51 0.552 0.584 0.522 ...
##  $ process.batch : int  1 1 1 1 1 1 1 1 1 1 ...
##  $ Data.File     : chr  "1.D" "2.D" "3.D" "4.D" ...
##  $ Type          : chr  "Sample" "Sample" "Sample" "Sample" ...
##  $ Acq..Date.Time: chr  "18/08/2022 23:28" "19/08/2022 00:19" "19/08/2022 01:11" "19/08/2022 02:02" ...
##  $ C1            : num  0.0144 0.0172 0.0477 0.0151 0.0411 ...
##  $ C2            : num  0.00387 0.00186 0.0089 0.00493 0.00455 ...
##  $ C3            : num  0.3128 0.2537 0.0413 0.1989 0.299 ...
##  $ C4            : num  0.0233 0.0283 0.0718 0.0375 0.053 ...
##  $ C5            : num  0.0333 0.037 0.1111 0.0556 0.0703 ...
##  $ C6            : num  0.506 0.259 0.779 0.539 0.481 ...
##  $ C7            : num  0.0638 0.0185 0.1387 0.0411 0.0427 ...
##  $ C8            : num  0.283 0.193 0.662 0.373 0.377 ...
##  $ C9            : num  0.01558 0.00917 0.0323 0.01178 0.0137 ...
##  $ C10           : num  0.00936 0.00455 0.01574 0.00549 0.00751 ...
##  $ C11           : num  0.00436 0.00299 0.01354 0.00311 0.00504 ...
##  $ C12           : num  0.00933 0.00623 0.02246 0.01486 0.01038 ...
##  $ C13           : num  0.02815 0.00293 0.10157 0.06982 0.00502 ...
##  $ C14           : num  0.0734 0.0186 0.0904 0.0498 0.0275 ...
##  $ C15           : num  0.44 0.155 0.619 0.132 0.256 ...
##  $ C16           : num  0.00557 0.00436 0.00646 0.00033 0.00962 ...
##  $ C17           : num  0.0564 0.0275 0.1053 0.1022 0.05 ...
##  $ C18           : num  0.1165 0.0536 0.2856 0.1701 0.1246 ...
##  $ C19           : num  0.00626 0.04157 0.01157 0.00132 0.06392 ...
##  $ C20           : num  0.0365 0.0331 0.0471 0.056 0.0512 ...
##  $ C21           : num  0.1211 0.0875 0.2304 0.1267 0.1918 ...
```

```
CHC$Mass2    <- CHC$Mass * 0.75
CHC$mito.nuc <- interaction(CHC$Nuclear, CHC$Mito, drop = TRUE)

man.response <- as.matrix(CHC.cor)
```

---

### 5 Compositional log-ratio transformation and covariate adjustment

Because CHCs form compositional data (each compound is a proportion
of the total blend), raw values are first closed to proportions, then
transformed using the centred log-ratio (CLR) transform. Importantly,
covariate adjustment for experimental block and allometric body mass
(Mass^(2/3), reflecting surface-area scaling) is performed **in
CLR space** after transformation. This preserves Aitchison
geometry and ensures that distance-based variance partitioning is
conducted in a geometrically consistent space.

```
# 1) Closure: convert to proportions
P  <- as.matrix(CHC.cor)
rs <- rowSums(P, na.rm = TRUE)
keep <- is.finite(rs) & (rs > 0)
P  <- P[keep, , drop = FALSE]
rs <- rs[keep]
P  <- P / rs

# 2) Treat very small values as zeros for replacement
tol     <- .Machine$double.eps^0.5   # ~1.5e-8
P[P < tol] <- 0

# 3) Multiplicative zero replacement if needed
if (any(P == 0, na.rm = TRUE)) {
  P_nz <- zCompositions::cmultRepl(P, label = 0, method = "CZM")
} else {
  P_nz <- P
}

# 4) CLR transform
Y_clr <- as.matrix(compositions::clr(acomp(P_nz)))

# 5) Subset metadata to match rows retained after closure
meta_keep <- CHC[keep, , drop = FALSE]

# 6) Regress out block and allometric mass in CLR space
blk         <- factor(meta_keep$block)
mass_cov    <- I(meta_keep$Mass^(2/3))
Y_clr_resid <- resid(lm(Y_clr ~ blk + mass_cov))
```

---

### 6 Align metadata

```
# meta_keep was built from CHC[keep, ] above, so rows are already aligned
meta_clr         <- meta_keep
meta_clr$Nuclear <- droplevels(factor(meta_clr$Nuclear))
meta_clr$Mito    <- droplevels(factor(meta_clr$Mito))
meta_clr$block   <- droplevels(factor(meta_clr$block))
```

---

### 7 Shared colour palette

All figures in this pipeline use a fixed, colourblind-safe palette
based on Okabe & Ito (2008). The same genotype colours are applied
consistently across the PCA and heatmap.

```
# Genotype palette — ggplot hue_pal() with "i" swapped to clear purple.
# Matches the palette used in the main paper figures.
geno_levels <- c("A", "B", "C", "D", "E", "F", "G", "H", "i")

geno_cols        <- setNames(hue_pal()(9), geno_levels)
geno_cols["i"]   <- "#7B4F9E"  # replace near-default-pink with clear purple

nuc_cols  <- geno_cols
mito_cols <- geno_cols

# Mating category colours (consistent with Pipelines 2 & 3)
type_cols <- c(
  "Match"     = "#E76F51",  # co-evolved — warm terracotta
  "NucMatch"  = "#457B9D",  # nuclear match only — steel blue
  "MitoMatch" = "#2A9D8F"   # mito match only — teal
)
```

---

### 8 Principal component analysis

PCA is performed on the CLR-transformed, covariate-adjusted
residuals. Panel A shows the overall separation of nuclear genotypes;
Panel B facets by mitochondrial haplotype to reveal how mtDNA shifts
genotype clusters, consistent with a mitonuclear interaction.

```
stopifnot(nrow(Y_clr_resid) == nrow(meta_clr))
Y <- as.matrix(Y_clr_resid)

pca    <- prcomp(Y, center = TRUE, scale. = FALSE)
var_pc <- round(100 * summary(pca)$importance[2, 1:5], 1)
var_pc
```

```
##  PC1  PC2  PC3  PC4  PC5 
## 44.8 15.2 12.8  8.2  4.5
```

```
# Scores + metadata
pcadf <- data.frame(
  PC1     = pca$x[, 1],
  PC2     = pca$x[, 2],
  PC3     = pca$x[, 3],
  PC4     = pca$x[, 4],
  PC5     = pca$x[, 5],
  Nuclear = droplevels(meta_clr$Nuclear),
  Mito    = droplevels(meta_clr$Mito)
)

# Colours defined in the shared palette chunk above

# Panel A: overall by nuclear background
pCLR1 <- ggplot(pcadf, aes(PC1, PC2, colour = Nuclear)) +
  geom_point(alpha = 0.6, size = 2.2) +
  stat_ellipse(aes(group = Nuclear), level = 0.68, linewidth = 0.9) +
  labs(x = sprintf("PC1 (%.1f%%)", var_pc[1]),
       y = sprintf("PC2 (%.1f%%)", var_pc[2])) +
  scale_color_manual(values = nuc_cols) +
  theme_bw() +
  theme(legend.position = "right")

# Panel B: faceted by mitochondrial haplotype
pCLR2 <- ggplot(pcadf, aes(PC1, PC2, colour = Nuclear)) +
  geom_point(alpha = 0.6, size = 2.0) +
  stat_ellipse(aes(group = Nuclear), level = 0.68, linewidth = 0.8) +
  labs(x = sprintf("PC1 (%.1f%%)", var_pc[1]),
       y = sprintf("PC2 (%.1f%%)", var_pc[2])) +
  facet_wrap(~ Mito, ncol = 3,
             labeller = labeller(Mito = function(x) paste("mtDNA:", x))) +
  scale_color_manual(values = nuc_cols) +
  theme_bw() +
  theme(strip.background = element_rect(fill = "grey90"),
        strip.text       = element_text(face = "bold"),
        legend.position  = "none")

fig_pca <- (pCLR1 + pCLR2) +
  plot_annotation(tag_levels = "A") &
  theme(plot.tag = element_text(face = "bold", size = 18))

fig_pca
```

---

### 9 PERMANOVA

Distance-based multivariate analysis of variance (PERMANOVA; 9,999
permutations) partitions Euclidean distances in CLR space among nuclear
background, mitochondrial haplotype, and their interaction.

```
D_clr <- dist(Y_clr_resid, method = "euclidean")

# Marginal main effects
adonis_main_clr <- adonis2(D_clr ~ Nuclear + Mito, data = meta_clr,
                           by = "margin", permutations = 9999)
print(adonis_main_clr)
```

```
## Permutation test for adonis under reduced model
## Marginal effects of terms
## Permutation: free
## Number of permutations: 9999
## 
## adonis2(formula = D_clr ~ Nuclear + Mito, data = meta_clr, permutations = 9999, by = "margin")
##           Df SumOfSqs      R2      F Pr(>F)    
## Nuclear    8  1610.11 0.57016 93.849  1e-04 ***
## Mito       8    47.95 0.01698  2.795  1e-04 ***
## Residual 538  1153.77 0.40856                  
## Total    554  2823.97 1.00000                  
## ---
## Signif. codes:  0 '***' 0.001 '**' 0.01 '*' 0.05 '.' 0.1 ' ' 1
```

```
# Marginal interaction term (Type-III-like)
adonis_int_clr <- adonis2(D_clr ~ Nuclear + Mito + Nuclear:Mito,
                          data = meta_clr, by = "margin",
                          permutations = 9999)
print(adonis_int_clr)
```

```
## Permutation test for adonis under reduced model
## Marginal effects of terms
## Permutation: free
## Number of permutations: 9999
## 
## adonis2(formula = D_clr ~ Nuclear + Mito + Nuclear:Mito, data = meta_clr, permutations = 9999, by = "margin")
##               Df SumOfSqs      R2      F Pr(>F)    
## Nuclear:Mito  63   248.18 0.08788 2.0663  1e-04 ***
## Residual     475   905.58 0.32068                  
## Total        554  2823.97 1.00000                  
## ---
## Signif. codes:  0 '***' 0.001 '**' 0.01 '*' 0.05 '.' 0.1 ' ' 1
```

---

### 10 Dispersion diagnostic

PERMDISP tests whether the PERMANOVA results reflect differences in
group centroids (the intended interpretation) rather than differences in
within-group spread. A non-significant result supports a location rather
than dispersion interpretation.

```
grp_all <- interaction(meta_clr$Nuclear, meta_clr$Mito, drop = TRUE)
tab_g   <- table(grp_all)
keep_bd <- tab_g[grp_all] >= 2

if (sum(keep_bd) >= 2 && length(unique(grp_all[keep_bd])) >= 2) {
  grp_bd <- droplevels(grp_all[keep_bd])
  Y_bd   <- Y_clr_resid[keep_bd, , drop = FALSE]
  D_bd   <- dist(Y_bd, method = "euclidean")
  bd     <- betadisper(D_bd, grp_bd)

  print(anova(bd))
  set.seed(params$seed)
  print(permutest(bd, permutations = 9999))
} else {
  message("PERMDISP skipped: need ≥2 groups with ≥2 samples each.")
}
```

```
## Analysis of Variance Table
## 
## Response: Distances
##            Df  Sum Sq Mean Sq F value Pr(>F)
## Groups     79  26.522 0.33572  1.0665  0.338
## Residuals 475 149.523 0.31478               
## 
## Permutation test for homogeneity of multivariate dispersions
## Permutation: free
## Number of permutations: 9999
## 
## Response: Distances
##            Df  Sum Sq Mean Sq      F N.Perm Pr(>F)
## Groups     79  26.522 0.33572 1.0665   9999 0.3479
## Residuals 475 149.523 0.31478
```

---

### 11 RRPP MANOVA

MANOVA with residual randomisation permutation procedures (RRPP)
provides a location-focused test of genotype effects that is less
sensitive to dispersion heterogeneity than PERMANOVA. Nuclear
background, mitochondrial haplotype, and their interaction all produce
significant location shifts in mean CHC blend composition.

```
stopifnot(nrow(Y_clr_resid) == nrow(meta_clr))

Yc <- as.matrix(Y_clr_resid)
storage.mode(Yc) <- "double"
rownames(Yc)     <- rownames(meta_clr)

fit_clr <- procD.lm(Yc ~ Nuclear * Mito, data = meta_clr,
                    iter = 9999, SS.type = "III")
anova(fit_clr)
```

```
## 
## Analysis of Variance, using Residual Randomization
## Permutation procedure: Randomization of null model residuals 
## Number of permutations: 10000 
## Estimation method: Ordinary Least Squares 
## Sums of Squares and Cross-products: Type III 
## Effect sizes (Z) based on F distributions
## 
##               Df      SS     MS     Rsq       F       Z Pr(>F)    
## Nuclear        8  255.91 31.988 0.09062 16.7786 11.8110  1e-04 ***
## Mito           8   58.53  7.316 0.02073  3.8374  6.8455  1e-04 ***
## Nuclear:Mito  63  248.18  3.939 0.08788  2.0663 10.1926  1e-04 ***
## Residuals    475  905.58  1.906 0.32068                           
## Total        554 2823.97                                          
## ---
## Signif. codes:  0 '***' 0.001 '**' 0.01 '*' 0.05 '.' 0.1 ' ' 1
## 
## Call: procD.lm(f1 = Yc ~ Nuclear * Mito, iter = 9999, SS.type = "III",  
##     data = meta_clr)
```

---

### 12 Which CHCs drive the mitonuclear interaction?

Species scores from an interaction-constrained ordination (partial
dbRDA, conditioning on nuclear and mitochondrial main effects) identify
which individual CHCs load most strongly on the unique mitonuclear
interaction axis in CLR space.

```
mod_I_clr <- capscale(
  Y_clr_resid ~ Nuclear:Mito + Condition(Nuclear) + Condition(Mito),
  data = meta_clr, distance = "euclidean"
)

sp_clr   <- scores(mod_I_clr, display = "species", choices = 1:2, scaling = 2)
int_rank <- sort(rowSums(sp_clr^2), decreasing = TRUE)
head(int_rank, 10)
```

```
##        C19        C16        C13         C3        C15         C2         C6 
## 0.96201959 0.59709715 0.35541803 0.15852553 0.11444006 0.08669273 0.05317666 
##         C7         C1        C17 
## 0.05022531 0.04210657 0.03960921
```

C19 is the strongest driver of the unique mitonuclear interaction,
followed by C16 (then C13, C15, and others). These compounds best
separate genotypes along the axis that is specifically attributable to
the interaction between nuclear and mitochondrial genomes.

---

### 13 Focal compound plots: C19 (heptacosene) and C16 (pentacosene)

C19 and C16 are the two compounds that load most strongly on the
unique mitonuclear interaction axis. Rather than summarising their
variation through ordination scores, these plots show their CLR-adjusted
values directly, averaged per mitonuclear genotype, arranged as a
nuclear × mito grid. This makes the interaction structure visually
explicit: if the compounds varied only with nuclear background, each row
would be flat; if only with mtDNA, each column would be flat. Departures
from both patterns indicate a genuine mitonuclear interaction.

```
# Identify columns — positional fallback if not named C19/C16
c19_col <- if ("C19" %in% colnames(Y_clr_resid)) "C19" else colnames(Y_clr_resid)[19]
c16_col <- if ("C16" %in% colnames(Y_clr_resid)) "C16" else colnames(Y_clr_resid)[16]

# Build flat data frame — completely explicit, no pipes
focal_raw <- data.frame(
  Nuclear  = as.character(meta_clr$Nuclear),
  Mito     = as.character(meta_clr$Mito),
  C19_val  = as.numeric(Y_clr_resid[, c19_col]),
  C16_val  = as.numeric(Y_clr_resid[, c16_col]),
  stringsAsFactors = FALSE
)

# Aggregate to genotype means using base R — avoids any dplyr masking issues
focal_means <- aggregate(
  cbind(C19_val, C16_val) ~ Nuclear + Mito,
  data = focal_raw,
  FUN  = mean,
  na.rm = TRUE
)

# Pivot to long form manually
focal_long_c19 <- data.frame(
  Nuclear  = focal_means$Nuclear,
  Mito     = focal_means$Mito,
  Compound = paste0(c19_col, ": Heptacos-X-ene (27:1)"),
  CLR_mean = focal_means$C19_val,
  stringsAsFactors = FALSE
)
focal_long_c16 <- data.frame(
  Nuclear  = focal_means$Nuclear,
  Mito     = focal_means$Mito,
  Compound = paste0(c16_col, ": Z-12-Pentacosene (3) (25:1)"),
  CLR_mean = focal_means$C16_val,
  stringsAsFactors = FALSE
)
focal_df <- rbind(focal_long_c19, focal_long_c16)
focal_df$Compound <- factor(focal_df$Compound,
                             levels = c(paste0(c19_col, ": Heptacos-X-ene (27:1)"),
                                        paste0(c16_col, ": Z-12-Pentacosene (3) (25:1)")))

cat("Columns:", paste(names(focal_df), collapse = ", "), "\n")
```

```
## Columns: Nuclear, Mito, Compound, CLR_mean
```

```
cat("Rows:", nrow(focal_df), "\n")
```

```
## Rows: 160
```

```
# Tile plot
p_tile <- ggplot(focal_df,
                 aes(x = Nuclear, y = Mito, fill = CLR_mean)) +
  geom_tile(colour = "white", linewidth = 0.5) +
  geom_text(aes(label = round(CLR_mean, 2)),
            size = 2.8, colour = "grey20") +
  facet_wrap(~ Compound, ncol = 1) +
  scale_fill_gradient2(
    low      = "#2166AC",
    mid      = "white",
    high     = "#D6604D",
    midpoint = 0,
    name     = "Mean CLR\n(block & mass\nadjusted)"
  ) +
  labs(
    x        = "Nuclear background",
    y        = "Mitochondrial haplotype",
    title    = "Mitonuclear interaction in key CHC compounds",
    subtitle = "Mean CLR-adjusted values per genotype combination"
  ) +
  theme_bw(base_size = 11) +
  theme(
    strip.background = element_rect(fill = "grey92"),
    strip.text       = element_text(face = "bold", size = 11),
    panel.grid       = element_blank(),
    legend.position  = "right"
  )

# Insert NA rows for the missing FnucxAmito genotype so the line
# for mtDNA A breaks between nuclear E and G rather than connecting them
fa_focal <- data.frame(
  Nuclear  = "F",
  Mito     = "A",
  Compound = levels(focal_df$Compound),
  CLR_mean = NA_real_,
  stringsAsFactors = FALSE
)
fa_focal$Compound <- factor(fa_focal$Compound, levels = levels(focal_df$Compound))
focal_df <- rbind(focal_df, fa_focal)

# Line plot
p_line <- ggplot(focal_df,
                 aes(x = Nuclear, y = CLR_mean,
                     colour = Mito, group = Mito)) +
  geom_line(linewidth = 0.9, alpha = 0.85) +
  geom_point(size = 2.2, alpha = 0.85) +
  facet_wrap(~ Compound, ncol = 1, scales = "free_y") +
  scale_colour_manual(values = mito_cols, name = "mtDNA haplotype") +
  labs(
    x        = "Nuclear background",
    y        = "Mean CLR (block & mass adjusted)",
    title    = "Interaction profiles: C19 and C16",
    subtitle = "Each line is one mtDNA haplotype across nuclear backgrounds"
  ) +
  theme_bw(base_size = 11) +
  theme(
    strip.background = element_rect(fill = "grey92"),
    strip.text       = element_text(face = "bold", size = 11),
    panel.grid.minor = element_blank(),
    legend.position  = "right"
  )

fig_focal <- p_tile | p_line
fig_focal +
  plot_annotation(tag_levels = "A") &
  theme(plot.tag = element_text(face = "bold", size = 14))
```

```
ggsave(
  file.path(out_dir, "FigureS_focal_compounds.pdf"),
  plot   = fig_focal + plot_annotation(tag_levels = "A") &
           theme(plot.tag = element_text(face = "bold", size = 14)),
  device = "pdf", width = 14, height = 8, units = "in", dpi = 300
)
message("Saved: FigureS_focal_compounds.pdf")
```

---

### 14 All-compound interaction profiles

The same interaction line plot extended to all 21 CHC compounds,
sorted by their loading on the mitonuclear interaction axis (strongest
top-left). Non-parallel lines indicate a compound-specific mitonuclear
interaction; parallel lines indicate additive nuclear + mitochondrial
effects only. C19 and C16 appear first as they are the strongest
interaction drivers.

```
# Compound name lookup — from Supplementary Table S2
chc_names <- c(
  C1  = "C1: Heneicosane (21:0)",
  C2  = "C2: 1-Docosene (22:1)",
  C3  = "C3: Z-13-Octadecen-1-yl acetate (21:1 ester)",
  C4  = "C4: Docosane (22:0)",
  C5  = "C5: 9-Tricosene Z (1) (23:1)",
  C6  = "C6: 9-Tricosene Z (2) (23:1)",
  C7  = "C7: 9-Tricosene Z (3) (23:1)",
  C8  = "C8: Tricosane (23:0)",
  C9  = "C9: Tetracos-X-ene (1) (24:1)",
  C10 = "C10: Tetracos-X-ene (2) (24:1)",
  C11 = "C11: Tetracose-X-ene (3) (24:1)",
  C12 = "C12: Tetracosane (24:0)",
  C13 = "C13: 2-Methyltetracosane (25:0 br)",
  C14 = "C14: Z-12-Pentacosene (1) (25:1)",
  C15 = "C15: Z-12-Pentacosene (2) (25:1)",
  C16 = "C16: Z-12-Pentacosene (3) (25:1)",
  C17 = "C17: Pentacosane (25:0)",
  C18 = "C18: 2-Methylhexacosane (27:0 br)",
  C19 = "C19: Heptacos-X-ene (27:1)",
  C20 = "C20: Heptacosane (27:0)",
  C21 = "C21: 2-Methyloctacosane (29:0 br)"
)

# Fall back gracefully: if a compound name isn't in the lookup, use its column name
label_compound <- function(col_name, rank_i) {
  chem <- if (col_name %in% names(chc_names)) chc_names[col_name] else col_name
  paste0(chem, "\n(interaction rank ", rank_i, ")")
}

# Use int_rank (computed in chc-interaction-drivers) to order compounds
compound_order <- names(int_rank)

# Build genotype-mean CLR data for all compounds using base R
all_raw <- data.frame(
  Nuclear = as.character(meta_clr$Nuclear),
  Mito    = as.character(meta_clr$Mito),
  as.data.frame(Y_clr_resid),
  stringsAsFactors = FALSE
)

all_means <- aggregate(
  . ~ Nuclear + Mito,
  data = all_raw,
  FUN  = mean,
  na.rm = TRUE
)

# Pivot to long form using base R reshape
all_long <- reshape(
  all_means,
  varying      = compound_order,
  v.names      = "CLR_mean",
  timevar      = "Compound",
  times        = compound_order,
  direction    = "long"
)
all_long$Compound <- factor(all_long$Compound, levels = compound_order)
all_long$id       <- NULL

# Insert NA rows for the missing FnucxAmito (FA) genotype so geom_line
# breaks between E and G for the A mtDNA line rather than connecting them
fa_na <- data.frame(
  Nuclear   = "F",
  Mito      = "A",
  CLR_mean  = NA_real_,
  Compound  = factor(compound_order, levels = compound_order),
  stringsAsFactors = FALSE
)
all_long <- rbind(all_long, fa_na)

# Build facet labels: chemical name + interaction rank
rank_labels <- setNames(
  mapply(label_compound, names(int_rank), seq_along(int_rank)),
  names(int_rank)
)
all_long$CompoundLabel <- rank_labels[as.character(all_long$Compound)]
all_long$CompoundLabel <- factor(all_long$CompoundLabel,
                                  levels = rank_labels[compound_order])

fig_all <- ggplot(all_long,
                  aes(x = Nuclear, y = CLR_mean,
                      colour = Mito, group = Mito)) +
  geom_line(linewidth = 0.7, alpha = 0.8) +
  geom_point(size = 1.5, alpha = 0.8) +
  facet_wrap(~ CompoundLabel, ncol = 3, scales = "free_y") +
  scale_colour_manual(values = mito_cols, name = "mtDNA haplotype",
                      guide  = guide_legend(nrow = 1)) +
  labs(
    x        = "Nuclear background",
    y        = "Mean CLR (block & mass adjusted)",
    title    = "Mitonuclear interaction profiles — all 21 CHC compounds",
    subtitle = "Compounds ordered by loading on the mitonuclear interaction axis (strongest first)"
  ) +
  theme_bw(base_size = 10) +
  theme(
    strip.background  = element_rect(fill = "grey92"),
    strip.text        = element_text(face = "bold", size = 8),
    panel.grid.minor  = element_blank(),
    axis.text.x       = element_text(size = 7),
    legend.position   = "bottom",
    legend.key.width  = unit(2, "lines"),
    legend.spacing.x  = unit(0.5, "lines")
  )

fig_all
```

```
ggsave(
  file.path(out_dir, "FigureS_all_compounds.pdf"),
  plot   = fig_all,
  device = "pdf",
  width  = 14, height = 18, units = "in", dpi = 300
)
message("Saved: FigureS_all_compounds.pdf")
```

---

### 15 Heatmap of mean CHC profiles

Mean CLR-transformed, covariate-adjusted CHC profiles are shown for
each of the 80 mitonuclear genotype combinations. Rows are individual
CHC compounds (clustered by Ward’s method); columns are genotypes
ordered by nuclear background then mitochondrial haplotype.

```
stopifnot(nrow(Y_clr_resid) == nrow(meta_clr))
chc_vars <- colnames(Y_clr_resid)

df <- cbind(meta_clr[, c("Nuclear", "Mito")], as.data.frame(Y_clr_resid)) %>%
  dplyr::mutate(mito.nuc = interaction(Nuclear, Mito, drop = TRUE))

means <- df %>%
  dplyr::group_by(mito.nuc, Nuclear, Mito) %>%
  dplyr::summarise(
    dplyr::across(all_of(chc_vars), ~ mean(.x, na.rm = TRUE)),
    .groups = "drop"
  ) %>%
  as.data.frame()

mat           <- as.matrix(means[, chc_vars])
rownames(mat) <- means$mito.nuc
mat_z         <- scale(mat, center = TRUE, scale = TRUE)

ann_col           <- data.frame(Nuclear = means$Nuclear, Mito = means$Mito)
rownames(ann_col) <- rownames(mat_z)
ord               <- order(means$Nuclear, means$Mito)

pheatmap(
  t(mat_z)[, ord],
  annotation_col    = ann_col[ord, , drop = FALSE],
  annotation_colors = list(Nuclear = nuc_cols, Mito = mito_cols),
  clustering_method = "ward.D2",
  show_colnames     = FALSE,
  fontsize_row      = 7
)
```

---

### 16 Export manuscript figures to PDF

This chunk saves each main manuscript figure as a high-resolution PDF
suitable for journal submission. Files are written to the working
directory.

```
# Set output directory — change this path as needed
out_dir <- "~/OneDrive - University College London/-Projects-Papers/Tom - CHC/figures"
dir.create(out_dir, showWarnings = FALSE, recursive = TRUE)

export_fig <- function(plot_obj, filename, width = 10, height = 7) {
  path <- file.path(out_dir, filename)
  ggsave(path, plot = plot_obj, device = "pdf",
         width = width, height = height, units = "in", dpi = 300)
  message("Saved: ", path)
}

# Figure 1: PCA
export_fig(fig_pca, "Figure1_PCA.pdf", width = 12, height = 5)

# Figure 2: Heatmap
# pheatmap does not return a standard ggplot object — save via pdf()
pdf(file.path(out_dir, "Figure2_Heatmap.pdf"), width = 10, height = 8)
pheatmap(
  t(mat_z)[, ord],
  annotation_col    = ann_col[ord, , drop = FALSE],
  annotation_colors = list(Nuclear = nuc_cols, Mito = mito_cols),
  clustering_method = "ward.D2",
  show_colnames     = FALSE,
  fontsize_row      = 7
)
dev.off()
message("Saved: ", file.path(out_dir, "Figure2_Heatmap.pdf"))
```

---

### 17 Session info

```
sessionInfo()
```

```
## R version 4.5.1 (2025-06-13)
## Platform: aarch64-apple-darwin20
## Running under: macOS Sequoia 15.6.1
## 
## Matrix products: default
## BLAS:   /Library/Frameworks/R.framework/Versions/4.5-arm64/Resources/lib/libRblas.0.dylib 
## LAPACK: /Library/Frameworks/R.framework/Versions/4.5-arm64/Resources/lib/libRlapack.dylib;  LAPACK version 3.12.1
## 
## locale:
## [1] en_US.UTF-8/en_US.UTF-8/en_US.UTF-8/C/en_US.UTF-8/en_US.UTF-8
## 
## time zone: Europe/London
## tzcode source: internal
## 
## attached base packages:
## [1] grid      stats     graphics  grDevices utils     datasets  methods  
## [8] base     
## 
## other attached packages:
##  [1] compositions_2.0-9    zCompositions_1.5.0-5 survival_3.8-3       
##  [4] truncnorm_1.0-9       MASS_7.3-65           patchwork_1.3.1      
##  [7] geomorph_4.0.10       Matrix_1.7-3          rgl_1.3.24           
## [10] RRPP_2.1.2            Rmisc_1.5.1           plyr_1.8.9           
## [13] lattice_0.22-7        ggrepel_0.9.6         pheatmap_1.0.13      
## [16] vegan_2.7-1           permute_0.9-8         tidyr_1.3.1          
## [19] dplyr_1.1.4           scales_1.4.0          ggplot2_3.5.2        
## 
## loaded via a namespace (and not attached):
##  [1] tensorA_0.36.2.1   sass_0.4.10        generics_0.1.4     robustbase_0.99-6 
##  [5] jpeg_0.1-11        digest_0.6.37      magrittr_2.0.3     bayesm_3.1-6      
##  [9] evaluate_1.0.4     RColorBrewer_1.1-3 fastmap_1.2.0      jsonlite_2.0.0    
## [13] ape_5.8-1          mgcv_1.9-3         purrr_1.0.4        jquerylib_0.1.4   
## [17] cli_3.6.5          rlang_1.1.6        splines_4.5.1      base64enc_0.1-3   
## [21] withr_3.0.2        cachem_1.1.0       yaml_2.3.10        tools_4.5.1       
## [25] parallel_4.5.1     vctrs_0.6.5        R6_2.6.1           lifecycle_1.0.4   
## [29] htmlwidgets_1.6.4  cluster_2.1.8.1    pkgconfig_2.0.3    pillar_1.11.0     
## [33] bslib_0.9.0        gtable_0.3.6       glue_1.8.0         Rcpp_1.1.0        
## [37] DEoptimR_1.1-4     xfun_0.52          tibble_3.3.0       tidyselect_1.2.1  
## [41] rstudioapi_0.17.1  knitr_1.50         farver_2.1.2       htmltools_0.5.8.1 
## [45] nlme_3.1-168       labeling_0.4.3     rmarkdown_2.29     compiler_4.5.1
```
