## Supplementary Code 2 for "Mitonuclear interactions shape male cuticular hydrocarbon profiles with consequences on mating success"

##### Supplementary Pipeline 2: Mating assay analysis

###### Tom Allison, Stuart Harrison, Nick Lane, M. Florencia Camus

#### 2026-03-08

- 1 Overview
- 2 Packages
- 3
  Shared colour palette
- 4 Load data
- 5 Copulation success
  - 5.1 Model
  - 5.2 Predicted probabilities
  - 5.3 Pairwise comparisons between
    mating categories
  - 5.4 Figure: predicted copulation
    probability by mating category
- 6 Latency to mating
  - 6.1 Overdispersion check
  - 6.2 Model with overdispersion
    correction
  - 6.3 Pairwise comparisons between
    mating categories
  - 6.4 Predicted latency by mating
    category
- 7 Summary figure
- 8 Export manuscript figures to
  PDF
- 9 Session info

### 1 Overview

This pipeline analyses mating assay data from the *Drosophila
melanogaster* mitonuclear panel. Two outcomes are examined: (1)
whether a male successfully mated within a 2-hour no-choice assay
(binary copulation success), and (2) latency to mating among successful
pairs. Males are classified into three categories — co-evolved (both
genomes from the same population), nuclear-match only, and
mitochondrial-match only — and compared using generalised linear mixed
models (GLMMs) with female nuclear background as a fixed effect and
experimental run nested within week as a random effect.

---

### 2 Packages

```
library(EnvStats)
library(Rmisc)
library(sjPlot)
library(lme4)
library(car)
library(GGally)
library(ggeffects)
library(emmeans)
library(performance)
library(ggplot2)
library(cowplot)
```

---

### 3 Shared colour palette

```
# Mating category colours — consistent across all three pipelines
type_cols <- c(
  "Match"     = "#E76F51",  # co-evolved — warm terracotta
  "NucMatch"  = "#457B9D",  # nuclear match only — steel blue
  "MitoMatch" = "#2A9D8F"   # mito match only — teal
)

type_labels <- c(
  "Match"     = "Co-evolved",
  "NucMatch"  = "Nuclear match only",
  "MitoMatch" = "Mitochondrial match only"
)
```

---

### 4 Load data

```
Mate <- read.csv(params$data_path, header = TRUE, strip.white = TRUE,
                 stringsAsFactors = FALSE)

Mate$Run     <- as.factor(Mate$Run)
Mate$Week    <- as.factor(Mate$Week)
Mate$Match   <- as.factor(Mate$Match)
Mate$week.run <- paste(Mate$Week, Mate$Run, sep = ".")

# Retain only the three mating categories used in this analysis
Mate <- subset(Mate, Type == "Match" | Type == "MitoMatch" | Type == "NucMatch")

head(Mate)
```

```
##   Female MaleNuc MaleMito Match      Type   TypeII Run Week Mating Latency Time
## 1      A       G        A     0 MitoMatch Mismatch   1    2      0      NA  120
## 2      A       A        A     1     Match    Match   1    2      0      NA  120
## 3      A       A        A     1     Match    Match   1    2      1      28   28
## 5      A       A        F     0  NucMatch Mismatch   1    2      0      NA  120
## 6      A       A        G     0  NucMatch Mismatch   1    2      1      90   90
## 7      B       H        B     0 MitoMatch Mismatch   1    2      0      NA  120
##   week.run
## 1      2.1
## 2      2.1
## 3      2.1
## 5      2.1
## 6      2.1
## 7      2.1
```

---

### 5 Copulation success

#### 5.1 Model

A binomial GLMM is used to model the binary outcome of whether a pair
mated within the 2-hour window. Mating category (`Type`) and
female nuclear background (`Female`) are included as fixed
effects; experimental run nested within week is included as a random
effect.

```
bin.mate <- glmer(
  Mating ~ Type + Female + (1 | Run %in% Week),
  data    = Mate,
  family  = "binomial",
  control = glmerControl(optimizer = "bobyqa", optCtrl = list(maxfun = 2e5))
)

Anova(bin.mate, type = 2)
```

```
## Analysis of Deviance Table (Type II Wald chisquare tests)
## 
## Response: Mating
##         Chisq Df Pr(>Chisq)    
## Type   22.712  2  1.170e-05 ***
## Female 52.253  8  1.504e-08 ***
## ---
## Signif. codes:  0 '***' 0.001 '**' 0.01 '*' 0.05 '.' 0.1 ' ' 1
```

```
summary(bin.mate)
```

```
## Generalized linear mixed model fit by maximum likelihood (Laplace
##   Approximation) [glmerMod]
##  Family: binomial  ( logit )
## Formula: Mating ~ Type + Female + (1 | Run %in% Week)
##    Data: Mate
## Control: glmerControl(optimizer = "bobyqa", optCtrl = list(maxfun = 2e+05))
## 
##       AIC       BIC    logLik -2*log(L)  df.resid 
##    1218.3    1276.9    -597.2    1194.3       960 
## 
## Scaled residuals: 
##     Min      1Q  Median      3Q     Max 
## -1.9472 -0.7549 -0.4842  0.9447  2.7781 
## 
## Random effects:
##  Groups   Name        Variance Std.Dev.
##  Run:Week (Intercept) 0.2728   0.5223  
## Number of obs: 972, groups:  Run:Week, 16
## 
## Fixed effects:
##               Estimate Std. Error z value Pr(>|z|)    
## (Intercept)    0.10058    0.28315   0.355 0.722423    
## TypeMitoMatch -0.85082    0.18971  -4.485 7.29e-06 ***
## TypeNucMatch  -0.57920    0.16824  -3.443 0.000576 ***
## FemaleB        0.54954    0.32100   1.712 0.086900 .  
## FemaleC        0.38272    0.30301   1.263 0.206565    
## FemaleD       -1.14828    0.31891  -3.601 0.000317 ***
## FemaleE       -0.20017    0.30138  -0.664 0.506581    
## FemaleF       -0.47526    0.33174  -1.433 0.151962    
## FemaleG       -0.06908    0.31203  -0.221 0.824783    
## FemaleH        0.30747    0.29738   1.034 0.301172    
## Femalei       -0.79146    0.34141  -2.318 0.020437 *  
## ---
## Signif. codes:  0 '***' 0.001 '**' 0.01 '*' 0.05 '.' 0.1 ' ' 1
## 
## Correlation of Fixed Effects:
##             (Intr) TypMtM TypNcM FemalB FemalC FemalD FemalE FemalF FemalG
## TypeMitMtch -0.313                                                        
## TypeNucMtch -0.308  0.446                                                 
## FemaleB     -0.568 -0.011 -0.005                                          
## FemaleC     -0.603  0.009 -0.004  0.532                                   
## FemaleD     -0.611  0.068  0.045  0.506  0.535                            
## FemaleE     -0.625  0.033  0.009  0.535  0.565  0.593                     
## FemaleF     -0.559  0.027  0.013  0.484  0.511  0.488  0.516              
## FemaleG     -0.599  0.019  0.017  0.521  0.548  0.526  0.554  0.502       
## FemaleH     -0.642  0.033  0.018  0.547  0.577  0.588  0.611  0.525  0.568
## Femalei     -0.559  0.068  0.038  0.471  0.499  0.485  0.506  0.454  0.486
##             FemalH
## TypeMitMtch       
## TypeNucMtch       
## FemaleB           
## FemaleC           
## FemaleD           
## FemaleE           
## FemaleF           
## FemaleG           
## FemaleH           
## Femalei      0.522
```

#### 5.2 Predicted probabilities

```
sjPlot::get_model_data(bin.mate, type = "pred")
```

```
## $Type
## # Predicted probabilities of Mating
## 
## Type | Predicted |     95% CI | group_col
## -----------------------------------------
##    1 |      0.53 | 0.39, 0.66 |      Type
##    2 |      0.32 | 0.21, 0.45 |      Type
##    3 |      0.38 | 0.26, 0.52 |      Type
## 
## Adjusted for:
## * Female = A
## 
## $Female
## # Predicted probabilities of Mating
## 
## Female | Predicted |     95% CI | group_col
## -------------------------------------------
##      1 |      0.53 | 0.39, 0.66 |    Female
##      2 |      0.66 | 0.52, 0.77 |    Female
##      3 |      0.62 | 0.49, 0.73 |    Female
##      4 |      0.26 | 0.17, 0.37 |    Female
##      5 |      0.48 | 0.36, 0.60 |    Female
##      6 |      0.41 | 0.28, 0.55 |    Female
##      7 |      0.51 | 0.38, 0.64 |    Female
##      9 |      0.33 | 0.22, 0.47 |    Female
## 
## Adjusted for:
## * Type = Match
```

#### 5.3 Pairwise comparisons between mating categories

```
test_predictions(bin.mate, "Type")
```

```
## # Pairwise comparisons
## 
## Type               | Contrast |       95% CI |      p
## -----------------------------------------------------
## Match-NucMatch     |     0.13 |  0.06,  0.21 | < .001
## MitoMatch-Match    |    -0.19 | -0.27, -0.11 | < .001
## MitoMatch-NucMatch |    -0.06 | -0.13,  0.02 | 0.149
```

#### 5.4 Figure: predicted copulation probability by mating category

```
dat <- predict_response(bin.mate, terms = c("Type"))

# Manual ggplot for consistent colour scheme
f1 <- ggplot(as.data.frame(dat),
             aes(x = x, y = predicted, colour = x, fill = x)) +
  geom_ribbon(aes(ymin = conf.low, ymax = conf.high), alpha = 0.2,
              colour = NA) +
  geom_point(size = 3) +
  geom_errorbar(aes(ymin = conf.low, ymax = conf.high),
                width = 0.15, linewidth = 0.7) +
  scale_colour_manual(values = type_cols, labels = type_labels,
                      name = NULL) +
  scale_fill_manual(values   = type_cols, labels = type_labels,
                    name = NULL) +
  scale_x_discrete(labels = type_labels) +
  labs(x = "Male genotype category",
       y = "Predicted copulation probability") +
  theme_bw(base_size = 11) +
  theme(legend.position = "none",
        panel.grid.minor = element_blank())
f1
```

---

### 6 Latency to mating

Analysis is restricted to pairs that successfully mated within the
observation window.

#### 6.1 Overdispersion check

Time-to-mating data follow a Poisson distribution. An initial model
is first fitted to check for overdispersion.

```
Mate.w <- subset(Mate, Mating == 1)

hist(Mate.w$Time, main = "Distribution of mating latency (minutes)",
     xlab = "Time to mating (min)")
```

```
p.model.init <- glmer(
  Time ~ Type + Female + (1 | Run %in% Week),
  data    = Mate.w,
  family  = "poisson",
  control = glmerControl(optimizer = "bobyqa", optCtrl = list(maxfun = 2e5))
)

check_overdispersion(p.model.init)
```

```
## # Overdispersion test
## 
##        dispersion ratio =   22.042
##   Pearson's Chi-Squared = 8067.495
##                 p-value =  < 0.001
```

#### 6.2 Model with overdispersion correction

An observation-level random effect is added to account for
overdispersion.

```
Mate.w$ID <- seq.int(nrow(Mate.w))

p.model <- glmer(
  Time ~ Type + Female + (1 | Run %in% Week) + (1 | ID),
  data    = Mate.w,
  family  = "poisson",
  control = glmerControl(optimizer = "bobyqa", optCtrl = list(maxfun = 2e5))
)

Anova(p.model, type = 2)
```

```
## Analysis of Deviance Table (Type II Wald chisquare tests)
## 
## Response: Time
##          Chisq Df Pr(>Chisq)    
## Type    1.8063  2     0.4053    
## Female 33.4363  8  5.138e-05 ***
## ---
## Signif. codes:  0 '***' 0.001 '**' 0.01 '*' 0.05 '.' 0.1 ' ' 1
```

```
summary(p.model)
```

```
## Generalized linear mixed model fit by maximum likelihood (Laplace
##   Approximation) [glmerMod]
##  Family: poisson  ( log )
## Formula: Time ~ Type + Female + (1 | Run %in% Week) + (1 | ID)
##    Data: Mate.w
## Control: glmerControl(optimizer = "bobyqa", optCtrl = list(maxfun = 2e+05))
## 
##       AIC       BIC    logLik -2*log(L)  df.resid 
##    3489.0    3540.1   -1731.5    3463.0       365 
## 
## Scaled residuals: 
##      Min       1Q   Median       3Q      Max 
## -1.38211 -0.21470  0.03325  0.10834  0.26887 
## 
## Random effects:
##  Groups   Name        Variance  Std.Dev.
##  ID       (Intercept) 0.7844024 0.88566 
##  Run:Week (Intercept) 0.0002749 0.01658 
## Number of obs: 378, groups:  ID, 378; Run:Week, 16
## 
## Fixed effects:
##                Estimate Std. Error z value Pr(>|z|)    
## (Intercept)    3.332941   0.166717  19.992  < 2e-16 ***
## TypeMitoMatch -0.172124   0.129060  -1.334  0.18231    
## TypeNucMatch  -0.067737   0.109656  -0.618  0.53676    
## FemaleB       -0.664063   0.208329  -3.188  0.00143 ** 
## FemaleC        0.362453   0.199279   1.819  0.06894 .  
## FemaleD       -0.084871   0.226565  -0.375  0.70796    
## FemaleE       -0.042139   0.202198  -0.208  0.83491    
## FemaleF        0.089779   0.236512   0.380  0.70424    
## FemaleG       -0.008561   0.213962  -0.040  0.96808    
## FemaleH        0.093818   0.190317   0.493  0.62204    
## Femalei       -0.112149   0.251959  -0.445  0.65624    
## ---
## Signif. codes:  0 '***' 0.001 '**' 0.01 '*' 0.05 '.' 0.1 ' ' 1
## 
## Correlation of Fixed Effects:
##             (Intr) TypMtM TypNcM FemalB FemalC FemalD FemalE FemalF FemalG
## TypeMitMtch -0.333                                                        
## TypeNucMtch -0.262  0.346                                                 
## FemaleB     -0.705  0.051 -0.024                                          
## FemaleC     -0.744  0.100 -0.032  0.589                                   
## FemaleD     -0.652  0.060 -0.033  0.525  0.545                            
## FemaleE     -0.735  0.077 -0.013  0.585  0.609  0.544                     
## FemaleF     -0.622  0.018  0.058  0.486  0.505  0.447  0.501              
## FemaleG     -0.674 -0.002 -0.030  0.553  0.570  0.513  0.570  0.474       
## FemaleH     -0.770  0.071 -0.021  0.612  0.641  0.566  0.632  0.529  0.593
## Femalei     -0.575  0.108 -0.052  0.445  0.482  0.408  0.460  0.391  0.420
##             FemalH
## TypeMitMtch       
## TypeNucMtch       
## FemaleB           
## FemaleC           
## FemaleD           
## FemaleE           
## FemaleF           
## FemaleG           
## FemaleH           
## Femalei      0.502
```

#### 6.3 Pairwise comparisons between mating categories

```
headcomps.p <- emmeans(p.model, "Type")
plot(headcomps.p, comparisons = TRUE)
```

#### 6.4 Predicted latency by mating category

```
sjPlot::get_model_data(p.model, type = "pred")
```

```
## $Type
## # Predicted counts of Time
## 
## Type | Predicted |       95% CI | group_col
## -------------------------------------------
##    1 |     28.02 | 20.21, 38.85 |      Type
##    2 |     23.59 | 16.96, 32.82 |      Type
##    3 |     26.19 | 18.63, 36.81 |      Type
## 
## Adjusted for:
## * Female = A
## 
## $Female
## # Predicted counts of Time
## 
## Female | Predicted |       95% CI | group_col
## ---------------------------------------------
##      1 |     28.02 | 20.21, 38.85 |    Female
##      2 |     14.42 | 10.79, 19.28 |    Female
##      3 |     40.26 | 30.94, 52.39 |    Female
##      4 |     25.74 | 18.40, 36.01 |    Female
##      5 |     26.86 | 20.54, 35.14 |    Female
##      6 |     30.65 | 21.29, 44.14 |    Female
##      7 |     27.78 | 20.44, 37.77 |    Female
##      9 |     25.05 | 16.88, 37.16 |    Female
## 
## Adjusted for:
## * Type = Match
```

---

### 7 Summary figure

Both outcomes — copulation success and mating latency — are shown
side by side.

```
dat2 <- predict_response(p.model, terms = c("Type"))

f2 <- ggplot(as.data.frame(dat2),
             aes(x = x, y = predicted, colour = x, fill = x)) +
  geom_ribbon(aes(ymin = conf.low, ymax = conf.high), alpha = 0.2,
              colour = NA) +
  geom_point(size = 3) +
  geom_errorbar(aes(ymin = conf.low, ymax = conf.high),
                width = 0.15, linewidth = 0.7) +
  scale_colour_manual(values = type_cols, labels = type_labels,
                      name = NULL) +
  scale_fill_manual(values   = type_cols, labels = type_labels,
                    name = NULL) +
  scale_x_discrete(labels = type_labels) +
  labs(x = "Male genotype category",
       y = "Predicted latency to mating (min)") +
  theme_bw(base_size = 11) +
  theme(legend.position = "none",
        panel.grid.minor = element_blank())

test_predictions(p.model, "Type")
```

```
## # Pairwise comparisons
## 
## Type               | Contrast |       95% CI |     p
## ----------------------------------------------------
## Match-MitoMatch    |     4.38 | -1.92, 10.69 | 0.173
## Match-NucMatch     |     1.82 | -3.92,  7.56 | 0.535
## NucMatch-MitoMatch |     2.57 | -4.01,  9.14 | 0.444
```

```
fig3 <- plot_grid(f1, f2, labels = c("A", "B"), align = "h")
fig3
```

---

### 8 Export manuscript figures to PDF

```
out_dir <- "~/OneDrive - University College London/-Projects-Papers/Tom - CHC/figures"
dir.create(out_dir, showWarnings = FALSE, recursive = TRUE)

# Figure 3: Mating summary (copulation success + latency)
ggsave(file.path(out_dir, "Figure3_Mating.pdf"),
       plot = fig3, device = "pdf",
       width = 10, height = 5, units = "in", dpi = 300)
message("Saved: Figure3_Mating.pdf")
```

---

### 9 Session info

```
sessionInfo()
```

```
## R version 4.5.1 (2025-06-13)
## Platform: aarch64-apple-darwin20
## Running under: macOS Sequoia 15.6.1
## 
## Matrix products: default
## BLAS:   /Library/Frameworks/R.framework/Versions/4.5-arm64/Resources/lib/libRblas.0.dylib 
## LAPACK: /Library/Frameworks/R.framework/Versions/4.5-arm64/Resources/lib/libRlapack.dylib;  LAPACK version 3.12.1
## 
## locale:
## [1] en_US.UTF-8/en_US.UTF-8/en_US.UTF-8/C/en_US.UTF-8/en_US.UTF-8
## 
## time zone: Europe/London
## tzcode source: internal
## 
## attached base packages:
## [1] stats     graphics  grDevices utils     datasets  methods   base     
## 
## other attached packages:
##  [1] cowplot_1.1.3      performance_0.15.1 emmeans_1.11.2     ggeffects_2.3.1   
##  [5] GGally_2.3.0       ggplot2_3.5.2      car_3.1-3          carData_3.0-5     
##  [9] lme4_1.1-37        Matrix_1.7-3       sjPlot_2.9.0       Rmisc_1.5.1       
## [13] plyr_1.8.9         lattice_0.22-7     EnvStats_3.1.0    
## 
## loaded via a namespace (and not attached):
##  [1] tidyr_1.3.1        sass_0.4.10        generics_0.1.4     digest_0.6.37     
##  [5] magrittr_2.0.3     estimability_1.5.1 evaluate_1.0.4     grid_4.5.1        
##  [9] RColorBrewer_1.1-3 mvtnorm_1.3-3      fastmap_1.2.0      jsonlite_2.0.0    
## [13] Formula_1.2-5      purrr_1.0.4        scales_1.4.0       jquerylib_0.1.4   
## [17] abind_1.4-8        reformulas_0.4.1   Rdpack_2.6.4       cli_3.6.5         
## [21] rlang_1.1.6        rbibutils_2.3      splines_4.5.1      withr_3.0.2       
## [25] cachem_1.1.0       yaml_2.3.10        datawizard_1.2.0   tools_4.5.1       
## [29] nloptr_2.2.1       minqa_1.2.8        dplyr_1.1.4        sjlabelled_1.2.0  
## [33] boot_1.3-31        ggstats_0.10.0     vctrs_0.6.5        R6_2.6.1          
## [37] lifecycle_1.0.4    MASS_7.3-65        insight_1.4.2      pkgconfig_2.0.3   
## [41] bslib_0.9.0        pillar_1.11.0      gtable_0.3.6       glue_1.8.0        
## [45] Rcpp_1.1.0         xfun_0.52          tibble_3.3.0       tidyselect_1.2.1  
## [49] rstudioapi_0.17.1  knitr_1.50         xtable_1.8-4       farver_2.1.2      
## [53] htmltools_0.5.8.1  nlme_3.1-168       labeling_0.4.3     rmarkdown_2.29    
## [57] compiler_4.5.1     S7_0.2.0
```
