## Supplementary Code 3 for "Mitonuclear interactions shape male cuticular hydrocarbon profiles with consequences on mating success"

##### Supplementary Pipeline 3: CHC axes predict copulation probability

###### Tom Allison, Stuart Harrison, Nick Lane, M. Florencia Camus

#### 2026-03-09

- 1 Overview
- 2 Packages
- 3
  Shared colour palette
- 4 Reconstruct CHC pipeline
  outputs
- 5 Compute per-genotype mean CHC PC
  scores
- 6 Load mating data and merge CHC
  scores
- 7 Logistic GLMM: CHC axes predicting
  copulation probability
  - 7.1 Model summary
  - 7.2 Type II ANOVA — CHC PC
    terms
  - 7.3 Likelihood ratio test: CHC axes vs
    null
- 8 Leave-one-genotype-out
  cross-validation (LOGO-CV)
- 9 Figures
- 10 Main paper figure
- 11 Export manuscript figures to
  PDF
- 12 Save results table
- 13 Session info

### 1 Overview

This analysis asks whether genotype-level CHC blend composition
predicts copulation probability across the mitonuclear panel. It bridges
the two existing pipelines — the CHC profiling pipeline
(`CHC-pipeline-log.rmd`) and the mating assay pipeline
(`mating_pipeline.rmd`) — by using genotype-mean PC scores
from CLR space as predictors in a logistic GLMM of mating outcome.

Two complementary analyses are performed:

1. **Logistic GLMM** — CHC PC axes as fixed-effect
   predictors of copulation success (binary), with female nuclear
   background and run-within-week as random effects (matching the original
   mating model structure).
2. **Leave-one-genotype-out cross-validation
   (LOGO-CV)** — for each of the 80 male genotypes in turn, the
   model is fit on all other genotypes and used to predict copulation
   probability for the held-out genotype. This provides an honest,
   out-of-sample estimate of how well CHC space predicts mating
   outcomes.

**Note on shared predictors:** Because CHC profiles were
measured at the genotype level (pooled triplets), all individuals of the
same male genotype share identical PC predictor values. The LOGO-CV
accounts for this by holding out entire genotypes rather than individual
observations.

---

### 2 Packages

```
library(dplyr)
library(tidyr)
library(lme4)
library(car)
library(ggplot2)
library(ggrepel)
library(patchwork)
library(zCompositions)
library(compositions)
```

---

### 3 Shared colour palette

```
# Mating category colours — consistent across all three pipelines
type_cols <- c(
  "Match"     = "#E76F51",  # co-evolved — warm terracotta
  "NucMatch"  = "#457B9D",  # nuclear match only — steel blue
  "MitoMatch" = "#2A9D8F"   # mito match only — teal
)

type_labels <- c(
  "Match"     = "Matched",
  "NucMatch"  = "Nuclear match only",
  "MitoMatch" = "Mitochondrial match only"
)
```

---

### 4 Reconstruct CHC pipeline outputs

This chunk re-runs the core CHC pipeline steps to produce
`pca` and `meta_clr`. If you have already run
`CHC-pipeline-log.rmd` in the same R session and these
objects are in your environment, you can skip this chunk by setting
`eval=FALSE` in the chunk header.

```
# Load raw CHC data
CHC.pre <- read.csv(params$chc_data_path, header = TRUE,
                    strip.white = TRUE, stringsAsFactors = FALSE)
CHC.pre <- CHC.pre[complete.cases(CHC.pre), ]
CHC.pre$Nuclear <- factor(CHC.pre$Nuclear)
CHC.pre$Mito    <- factor(CHC.pre$Mito)
CHC.pre$block   <- factor(CHC.pre$block)

# Internal standard correction
CHC.cor <- CHC.pre[, 13:33] / CHC.pre[, 12]
CHC     <- cbind(CHC.pre[, 1:10], CHC.cor)
CHC$mito.nuc <- interaction(CHC$Nuclear, CHC$Mito, drop = TRUE)

# Closure → zero replacement → CLR transform
P   <- as.matrix(CHC.cor)
rs  <- rowSums(P, na.rm = TRUE)
keep <- is.finite(rs) & (rs > 0)
P   <- P[keep, , drop = FALSE] / rs[keep]
tol <- .Machine$double.eps^0.5
P[P < tol] <- 0
P_nz <- if (any(P == 0)) zCompositions::cmultRepl(P, label = 0, method = "CZM") else P
Y_clr <- as.matrix(compositions::clr(acomp(P_nz)))

# Covariate adjustment in CLR space (block + allometric mass)
meta_keep   <- CHC[keep, , drop = FALSE]
blk         <- factor(meta_keep$block)
mass_cov    <- I(meta_keep$Mass^(2/3))
Y_clr_resid <- resid(lm(Y_clr ~ blk + mass_cov))

meta_clr         <- meta_keep
meta_clr$Nuclear <- droplevels(factor(meta_clr$Nuclear))
meta_clr$Mito    <- droplevels(factor(meta_clr$Mito))
meta_clr$block   <- droplevels(factor(meta_clr$block))

# PCA on CLR residuals
pca    <- prcomp(Y_clr_resid, center = TRUE, scale. = FALSE)
var_pc <- round(100 * summary(pca)$importance[2, 1:params$n_pcs], 1)
```

Variance explained by first 5 PCs: PC1 (44.8%), PC2 (15.2%), PC3
(12.8%), PC4 (8.2%), PC5 (4.5%).

---

### 5 Compute per-genotype mean CHC PC scores

```
pc_scores           <- as.data.frame(pca$x[, 1:n_pcs])
colnames(pc_scores) <- paste0("PC", 1:n_pcs)
pc_scores$Nuclear   <- meta_clr$Nuclear
pc_scores$Mito      <- meta_clr$Mito
pc_scores$mito.nuc  <- interaction(pc_scores$Nuclear, pc_scores$Mito,
                                    drop = TRUE)

geno_chc <- pc_scores %>%
  group_by(mito.nuc, Nuclear, Mito) %>%
  dplyr::summarise(across(starts_with("PC"), mean), .groups = "drop")

cat("CHC genotype means computed for", nrow(geno_chc), "genotypes\n")
```

```
## CHC genotype means computed for 80 genotypes
```

---

### 6 Load mating data and merge CHC scores

```
Mate <- read.csv(params$mate_data_path, header = TRUE,
                 strip.white = TRUE, stringsAsFactors = FALSE)

Mate$Run      <- factor(Mate$Run)
Mate$Week     <- factor(Mate$Week)
Mate$Female   <- factor(Mate$Female)
Mate$week.run <- paste(Mate$Week, Mate$Run, sep = ".")
Mate          <- subset(Mate, Type %in% c("Match", "MitoMatch", "NucMatch"))

# Build male genotype key from MaleNuc × MaleMito
Mate$MaleNuc  <- factor(Mate$MaleNuc)
Mate$MaleMito <- factor(Mate$MaleMito)
Mate$mito.nuc <- interaction(Mate$MaleNuc, Mate$MaleMito, drop = TRUE)

# Join genotype-mean PC scores
geno_chc_join <- geno_chc %>%
  mutate(mito.nuc = interaction(Nuclear, Mito, drop = TRUE))

Mate_chc <- Mate %>%
  left_join(dplyr::select(geno_chc_join, mito.nuc, starts_with("PC")),
            by = "mito.nuc")

n_missing <- sum(is.na(Mate_chc$PC1))
if (n_missing > 0) {
  warning(n_missing, " mating rows had no matching CHC genotype.")
  Mate_chc <- Mate_chc %>% filter(!is.na(PC1))
}

cat("Mating observations with CHC scores:", nrow(Mate_chc), "\n")
```

```
## Mating observations with CHC scores: 971
```

```
cat("Unique male genotypes:", length(unique(Mate_chc$mito.nuc)), "\n")
```

```
## Unique male genotypes: 80
```

---

### 7 Logistic GLMM: CHC axes predicting copulation probability

The model structure mirrors the original mating pipeline
(`bin.mate`): `Female` as a fixed effect and
`Run %in% Week` as a random effect. PC1–PC5 are added as
additional fixed-effect predictors. A likelihood ratio test against a
null model (female + random effects only) tests whether CHC axes
collectively explain variance in mating outcome.

```
pc_terms     <- paste(paste0("PC", 1:n_pcs), collapse = " + ")
full_formula <- as.formula(
  paste("Mating ~ Female +", pc_terms, "+ (1 | Run %in% Week)")
)
null_formula <- as.formula("Mating ~ Female + (1 | Run %in% Week)")

mod_full <- glmer(full_formula, data = Mate_chc,
                  family  = binomial(link = "logit"),
                  control = glmerControl(optimizer = "bobyqa",
                                         optCtrl   = list(maxfun = 2e5)))

mod_null <- glmer(null_formula, data = Mate_chc,
                  family  = binomial(link = "logit"),
                  control = glmerControl(optimizer = "bobyqa",
                                         optCtrl   = list(maxfun = 2e5)))
```

#### 7.1 Model summary

```
summary(mod_full)
```

```
## Generalized linear mixed model fit by maximum likelihood (Laplace
##   Approximation) [glmerMod]
##  Family: binomial  ( logit )
## Formula: Mating ~ Female + PC1 + PC2 + PC3 + PC4 + PC5 + (1 | Run %in%  
##     Week)
##    Data: Mate_chc
## Control: glmerControl(optimizer = "bobyqa", optCtrl = list(maxfun = 2e+05))
## 
##       AIC       BIC    logLik -2*log(L)  df.resid 
##    1208.9    1282.1    -589.5    1178.9       956 
## 
## Scaled residuals: 
##     Min      1Q  Median      3Q     Max 
## -2.1912 -0.7415 -0.4769  0.9019  4.0027 
## 
## Random effects:
##  Groups   Name        Variance Std.Dev.
##  Run:Week (Intercept) 0.2913   0.5397  
## Number of obs: 971, groups:  Run:Week, 16
## 
## Fixed effects:
##             Estimate Std. Error z value Pr(>|z|)    
## (Intercept)  0.18173    0.32122   0.566 0.571558    
## FemaleB     -0.03429    0.38626  -0.089 0.929256    
## FemaleC     -0.64926    0.37499  -1.731 0.083377 .  
## FemaleD     -1.23213    0.40559  -3.038 0.002383 ** 
## FemaleE     -0.99943    0.39155  -2.553 0.010695 *  
## FemaleF     -0.97226    0.41083  -2.367 0.017952 *  
## FemaleG     -0.66456    0.38584  -1.722 0.085005 .  
## FemaleH     -0.23792    0.36636  -0.649 0.516070    
## Femalei     -1.76294    0.43197  -4.081 4.48e-05 ***
## PC1          0.18002    0.07513   2.396 0.016574 *  
## PC2         -0.05423    0.12581  -0.431 0.666439    
## PC3          0.57519    0.15532   3.703 0.000213 ***
## PC4          0.66119    0.17414   3.797 0.000147 ***
## PC5          0.02259    0.26379   0.086 0.931750    
## ---
## Signif. codes:  0 '***' 0.001 '**' 0.01 '*' 0.05 '.' 0.1 ' ' 1
```

#### 7.2 Type II ANOVA — CHC PC terms

```
Anova(mod_full, type = 2)
```

```
## Analysis of Deviance Table (Type II Wald chisquare tests)
## 
## Response: Mating
##          Chisq Df Pr(>Chisq)    
## Female 39.3682  8    4.2e-06 ***
## PC1     5.7410  1  0.0165736 *  
## PC2     0.1858  1  0.6664385    
## PC3    13.7138  1  0.0002129 ***
## PC4    14.4160  1  0.0001465 ***
## PC5     0.0073  1  0.9317504    
## ---
## Signif. codes:  0 '***' 0.001 '**' 0.01 '*' 0.05 '.' 0.1 ' ' 1
```

#### 7.3 Likelihood ratio test: CHC axes vs null

```
anova(mod_null, mod_full)
```

```
## Data: Mate_chc
## Models:
## ..1: Mating ~ Female + (1 | Run %in% Week)
## ..2: Mating ~ Female + PC1 + PC2 + PC3 + PC4 + PC5 + (1 | Run %in% Week)
##     npar    AIC    BIC  logLik -2*log(L)  Chisq Df Pr(>Chisq)    
## ..1   10 1236.6 1285.4 -608.29    1216.6                         
## ..2   15 1208.9 1282.1 -589.46    1178.9 37.662  5  4.411e-07 ***
## ---
## Signif. codes:  0 '***' 0.001 '**' 0.01 '*' 0.05 '.' 0.1 ' ' 1
```

---

### 8 Leave-one-genotype-out cross-validation (LOGO-CV)

For each of the 81 male genotypes, the model is fit on all remaining
genotypes and used to predict copulation probability for the held-out
genotype (population-level prediction, marginalising over random
effects). Pearson *r* and Spearman ρ between predicted and
observed mating rates quantify out-of-sample predictive accuracy.

```
genotypes    <- levels(Mate_chc$mito.nuc)
logo_results <- vector("list", length(genotypes))

pb <- txtProgressBar(min = 0, max = length(genotypes), style = 3)

for (i in seq_along(genotypes)) {
  g     <- genotypes[i]
  train <- Mate_chc %>% filter(mito.nuc != g)
  test  <- Mate_chc %>% filter(mito.nuc == g)

  mod_cv <- tryCatch(
    suppressWarnings(
      glmer(full_formula, data = train, family = binomial(link = "logit"),
            control = glmerControl(optimizer = "bobyqa",
                                   optCtrl   = list(maxfun = 2e5)))
    ),
    error = function(e) NULL
  )

  if (!is.null(mod_cv)) {
    train_female_levels <- levels(droplevels(factor(train$Female)))
    test_pred <- test %>%
      mutate(Female = factor(Female, levels = train_female_levels)) %>%
      filter(!is.na(Female))

    pred_prob <- tryCatch({
      mean(predict(mod_cv, newdata = test_pred, type = "response",
                   re.form = NA, allow.new.levels = TRUE), na.rm = TRUE)
    }, error = function(e) NA_real_)

    obs_rate <- mean(test$Mating)
    n_obs    <- nrow(test)
  } else {
    pred_prob <- NA_real_
    obs_rate  <- mean(test$Mating)
    n_obs     <- nrow(test)
  }

  logo_results[[i]] <- data.frame(
    mito.nuc  = g,
    Nuclear   = as.character(unique(test$MaleNuc))[1],
    Mito      = as.character(unique(test$MaleMito))[1],
    obs_rate  = obs_rate,
    pred_prob = pred_prob,
    n_obs     = n_obs
  )

  setTxtProgressBar(pb, i)
}
close(pb)
```

```
logo_df       <- bind_rows(logo_results)
logo_complete <- logo_df %>% filter(!is.na(pred_prob))

cv_r   <- cor(logo_complete$obs_rate, logo_complete$pred_prob,
               method = "pearson")
cv_rho <- cor(logo_complete$obs_rate, logo_complete$pred_prob,
               method = "spearman")

# Rejoin mating category for plotting
geno_type <- Mate_chc %>%
  group_by(mito.nuc) %>%
  dplyr::summarise(Type = names(which.max(table(Type))), .groups = "drop")

logo_complete <- logo_complete %>%
  left_join(geno_type, by = "mito.nuc")

cat("Folds completed:", nrow(logo_complete), "/", length(genotypes), "\n")
```

```
## Folds completed: 80 / 81
```

```
cat("Pearson r (predicted vs observed): ", round(cv_r,   3), "\n")
```

```
## Pearson r (predicted vs observed):  0.429
```

```
cat("Spearman rho:                      ", round(cv_rho, 3), "\n")
```

```
## Spearman rho:                       0.452
```

---

### 9 Figures

```
# Colours defined in the shared palette chunk above

# Panel A: LOGO-CV predicted vs observed mating rate
p_cv <- ggplot(logo_complete,
               aes(x = pred_prob, y = obs_rate,
                   colour = Type, size = n_obs)) +
  geom_point(alpha = 0.8) +
  geom_abline(intercept = 0, slope = 1,
              linetype = "dashed", colour = "grey50") +
  geom_smooth(aes(group = 1), method = "lm", se = TRUE,
              colour = "black", linewidth = 0.7, alpha = 0.15) +
  annotate("text", x = Inf, y = -Inf, hjust = 1.1, vjust = -0.5,
           label = sprintf("Pearson r = %.2f\nSpearman \u03c1 = %.2f",
                           cv_r, cv_rho),
           size = 3.5) +
  scale_colour_manual(values = type_cols) +
  scale_size_continuous(range = c(2, 6), name = "N obs") +
  labs(x = "Predicted copulation probability (LOGO-CV)",
       y = "Observed mating rate",
       colour = "Mating category",
       title = "Cross-validated CHC prediction of copulation probability") +
  theme_bw()

# Panel B: Fixed-effect coefficients for PC terms
fe <- as.data.frame(summary(mod_full)$coefficients)
fe$term <- rownames(fe)
fe_pc <- fe %>%
  filter(grepl("^PC", term)) %>%
  mutate(term = factor(term, levels = rev(paste0("PC", 1:n_pcs))))

p_coef <- ggplot(fe_pc, aes(x = Estimate, y = term)) +
  geom_vline(xintercept = 0, linetype = "dashed", colour = "grey60") +
  geom_errorbarh(aes(xmin = Estimate - 1.96 * `Std. Error`,
                     xmax = Estimate + 1.96 * `Std. Error`),
                 height = 0.25, colour = "grey40") +
  geom_point(size = 3, colour = "#2A9D8F") +
  labs(x = "Log-odds coefficient (95% Wald CI)",
       y = "CHC axis",
       title = "Fixed-effect estimates") +
  theme_bw()

# Panels C - F : PC1 - PC5 vs observed mating rate
logo_pc <- logo_complete %>%
  left_join(dplyr::select(geno_chc, mito.nuc, PC1, PC2, PC3, PC4, PC5), by = "mito.nuc")

p_pc1 <- ggplot(logo_pc, aes(x = PC1, y = obs_rate, colour = Type)) +
  geom_point(alpha = 0.8, size = 2.5) +
  geom_smooth(aes(group = 1), method = "lm", se = TRUE,
              colour = "black", linewidth = 0.7, alpha = 0.15) +
  scale_colour_manual(values = type_cols) +
  labs(x = sprintf("PC1 (%.1f%% variance)", var_pc[1]),
       y = "Observed mating rate") +
  theme_bw() + theme(legend.position = "none")

p_pc2 <- ggplot(logo_pc, aes(x = PC2, y = obs_rate, colour = Type)) +
  geom_point(alpha = 0.8, size = 2.5) +
  geom_smooth(aes(group = 1), method = "lm", se = TRUE,
              colour = "black", linewidth = 0.7, alpha = 0.15) +
  scale_colour_manual(values = type_cols) +
  labs(x = sprintf("PC2 (%.1f%% variance)", var_pc[2]),
       y = "Observed mating rate") +
  theme_bw() + theme(legend.position = "none")

p_pc3 <- ggplot(logo_pc, aes(x = PC3, y = obs_rate, colour = Type)) +
  geom_point(alpha = 0.8, size = 2.5) +
  geom_smooth(aes(group = 1), method = "lm", se = TRUE,
              colour = "black", linewidth = 0.7, alpha = 0.15) +
  scale_colour_manual(values = type_cols) +
  labs(x = sprintf("PC3 (%.1f%% variance)", var_pc[3]),
       y = "Observed mating rate") +
  theme_bw() + theme(legend.position = "none")

p_pc4 <- ggplot(logo_pc, aes(x = PC4, y = obs_rate, colour = Type)) +
  geom_point(alpha = 0.8, size = 2.5) +
  geom_smooth(aes(group = 1), method = "lm", se = TRUE,
              colour = "black", linewidth = 0.7, alpha = 0.15) +
  scale_colour_manual(values = type_cols) +
  labs(x = sprintf("PC4 (%.1f%% variance)", var_pc[4]),
       y = "Observed mating rate") +
  theme_bw() + theme(legend.position = "none")

p_pc5 <- ggplot(logo_pc, aes(x = PC5, y = obs_rate, colour = Type)) +
  geom_point(alpha = 0.8, size = 2.5) +
  geom_smooth(aes(group = 1), method = "lm", se = TRUE,
              colour = "black", linewidth = 0.7, alpha = 0.15) +
  scale_colour_manual(values = type_cols) +
  labs(x = sprintf("PC5 (%.1f%% variance)", var_pc[5]),
       y = "Observed mating rate") +
  theme_bw() + theme(legend.position = "none")

(p_cv | p_coef) / (p_pc1 | p_pc2 | p_pc3 | p_pc4 | p_pc5 ) +
  plot_annotation(tag_levels = "A") &
  theme(plot.tag = element_text(face = "bold", size = 14))
```

---

### 10 Main paper figure

This is the cleaned-up figure intended for inclusion in the main
paper. It shows the cross-validated (LOGO-CV) predicted copulation
probability against the observed mating rate for each of the 80 male
genotypes, coloured by mating category. The nine co-evolved (Match)
genotypes are labelled directly. A horizontal reference line marks the
overall mean mating rate across all genotypes.

```
# Overall mean mating rate for reference line
mean_mating <- mean(Mate_chc$Mating)

# Flag co-evolved genotypes for labelling
logo_plot <- logo_complete %>%
  mutate(
    label     = ifelse(Type == "Match", as.character(mito.nuc), NA),
    coevolved = Type == "Match"
  )

# Colours and labels defined in the shared palette chunk above

# Panel B: LOGO-CV predicted vs observed mating rate
p_logo_main <- ggplot(logo_plot,
                   aes(x = pred_prob, y = obs_rate, colour = Type)) +
  # Reference lines
  geom_hline(yintercept = mean_mating,
             linetype = "dotted", colour = "grey60", linewidth = 0.6) +
  geom_abline(intercept = 0, slope = 1,
              linetype = "dashed", colour = "grey70", linewidth = 0.6) +
  # Points — co-evolved slightly larger and filled solid
  geom_point(aes(size = coevolved, alpha = coevolved)) +
  # Linear trend
  geom_smooth(aes(group = 1), method = "lm", se = TRUE,
              colour = "black", linewidth = 0.7, alpha = 0.12,
              show.legend = FALSE) +
  # Labels for co-evolved genotypes only
  geom_label_repel(aes(label = label),
                   size          = 2.8,
                   box.padding   = 0.3,
                   label.padding = 0.15,
                   label.size    = 0.2,
                   min.segment.length = 0.2,
                   show.legend   = FALSE,
                   na.rm         = TRUE) +
  # Correlation annotation
  annotate("text",
           x = 0.02, y = 0.97,
           hjust = 0, vjust = 1,
           label = sprintf("Pearson r = %.2f
Spearman ρ = %.2f",
                           cv_r, cv_rho),
           size = 3.2, colour = "grey30") +
  # Scales
  scale_colour_manual(values = type_cols, labels = type_labels,
                      name = "Male genotype") +
  scale_size_manual(values = c("FALSE" = 2.2, "TRUE" = 3.5),
                    guide = "none") +
  scale_alpha_manual(values = c("FALSE" = 0.65, "TRUE" = 0.95),
                     guide = "none") +
  scale_x_continuous(limits = c(0, 1), breaks = seq(0, 1, 0.2)) +
  scale_y_continuous(limits = c(0, 1), breaks = seq(0, 1, 0.2)) +
  # Labels
  labs(
    x = "Predicted copulation probability (LOGO cross-validation)",
    y = "Observed copulation probability"
  ) +
  theme_bw(base_size = 11) +
  theme(
    legend.position   = "right",
    legend.key.height = unit(0.9, "lines"),
    panel.grid.minor  = element_blank()
  )

# Main text figure: Panel A = fixed-effect coefficients, Panel B = LOGO-CV
fig_main <- (p_coef | p_logo_main) +
  plot_annotation(tag_levels = "A") &
  theme(plot.tag = element_text(face = "bold", size = 14))

fig_main
```

---

### 11 Export manuscript figures to PDF

```
out_dir <- "~/OneDrive - University College London/-Projects-Papers/Tom - CHC/figures"
dir.create(out_dir, showWarnings = FALSE, recursive = TRUE)

# Figure 4: CHC-mating prediction (main paper figure)
ggsave(file.path(out_dir, "Figure4_CHC_mating_prediction.pdf"),
       plot = fig_main, device = "pdf",
       width = 12, height = 5.5, units = "in", dpi = 300)
message("Saved: Figure4_CHC_mating_prediction.pdf")

# Supplementary: full diagnostic figure panel
ggsave(file.path(out_dir, "FigureS_CHC_mating_diagnostics.pdf"),
       plot = (p_cv | p_coef) / (p_pc1 | p_pc2) +
              plot_annotation(tag_levels = "A") &
              theme(plot.tag = element_text(face = "bold", size = 14)),
       device = "pdf", width = 12, height = 9, units = "in", dpi = 300)
message("Saved: FigureS_CHC_mating_diagnostics.pdf")
```

---

### 12 Save results table

```
logo_out <- logo_complete %>%
  left_join(geno_chc, by = c("mito.nuc", "Nuclear", "Mito")) %>%
  arrange(desc(obs_rate))

write.csv(logo_out, "CHC_mating_LOGO_results.csv", row.names = FALSE)
cat("Results saved to CHC_mating_LOGO_results.csv\n")
```

```
## Results saved to CHC_mating_LOGO_results.csv
```

---

### 13 Session info

```
sessionInfo()
```

```
## R version 4.5.1 (2025-06-13)
## Platform: aarch64-apple-darwin20
## Running under: macOS Sequoia 15.6.1
## 
## Matrix products: default
## BLAS:   /Library/Frameworks/R.framework/Versions/4.5-arm64/Resources/lib/libRblas.0.dylib 
## LAPACK: /Library/Frameworks/R.framework/Versions/4.5-arm64/Resources/lib/libRlapack.dylib;  LAPACK version 3.12.1
## 
## locale:
## [1] en_US.UTF-8/en_US.UTF-8/en_US.UTF-8/C/en_US.UTF-8/en_US.UTF-8
## 
## time zone: Europe/London
## tzcode source: internal
## 
## attached base packages:
## [1] stats     graphics  grDevices utils     datasets  methods   base     
## 
## other attached packages:
##  [1] compositions_2.0-9    zCompositions_1.5.0-5 survival_3.8-3       
##  [4] truncnorm_1.0-9       MASS_7.3-65           patchwork_1.3.1      
##  [7] ggrepel_0.9.6         ggplot2_3.5.2         car_3.1-3            
## [10] carData_3.0-5         lme4_1.1-37           Matrix_1.7-3         
## [13] tidyr_1.3.1           dplyr_1.1.4          
## 
## loaded via a namespace (and not attached):
##  [1] tensorA_0.36.2.1   sass_0.4.10        generics_0.1.4     robustbase_0.99-6 
##  [5] lattice_0.22-7     digest_0.6.37      magrittr_2.0.3     bayesm_3.1-6      
##  [9] evaluate_1.0.4     grid_4.5.1         RColorBrewer_1.1-3 fastmap_1.2.0     
## [13] jsonlite_2.0.0     Formula_1.2-5      mgcv_1.9-3         purrr_1.0.4       
## [17] scales_1.4.0       jquerylib_0.1.4    abind_1.4-8        reformulas_0.4.1  
## [21] Rdpack_2.6.4       cli_3.6.5          rlang_1.1.6        rbibutils_2.3     
## [25] splines_4.5.1      withr_3.0.2        cachem_1.1.0       yaml_2.3.10       
## [29] tools_4.5.1        nloptr_2.2.1       minqa_1.2.8        boot_1.3-31       
## [33] vctrs_0.6.5        R6_2.6.1           lifecycle_1.0.4    pkgconfig_2.0.3   
## [37] pillar_1.11.0      bslib_0.9.0        gtable_0.3.6       glue_1.8.0        
## [41] Rcpp_1.1.0         DEoptimR_1.1-4     xfun_0.52          tibble_3.3.0      
## [45] tidyselect_1.2.1   rstudioapi_0.17.1  knitr_1.50         farver_2.1.2      
## [49] htmltools_0.5.8.1  nlme_3.1-168       labeling_0.4.3     rmarkdown_2.29    
## [53] compiler_4.5.1
```
